# Identification of Dynamic Genetic Influences on DNA Methylation from Birth to Adulthood

**DOI:** 10.64898/2026.03.09.710470

**Authors:** Yueying Li, James R. Staley, Anna Großbach, Claudio Cappadona, Alexandre A. Lussier, Erin C. Dunn, Janine F. Felix, Charlotte A.M. Cecil, Dan J. Stein, Heather J. Zar, Venexia Walker, Tom R. Gaunt, Kate Tilling, Rosa Mulder, Andrew J. Simpkin, Gibran Hemani, Josine L. Min

## Abstract

Whether and how genetic regulation of DNA methylation (DNAm) change across the lifespan remains unclear. Here, we map age-dependent methylation quantitative trait loci (longitudinal mQTLs) using linear mixed models applied to repeated blood DNAm measures from birth, childhood, adolescence and adulthood in the Avon Longitudinal Study of Parents and Children. We identify 2,210 longitudinal mQTLs (2,393 SNP-CpG pairs; 7.3% *trans*) and observe consistent genotype-by-age effects in two independent cohorts of diverse ancestries (Pearson’s *r* = 0.85 in the Generation R Study; *r* = 0.56 in the Drakenstein Child Health Study). Longitudinal mQTLs show increasing effects with age at half of loci and associations with multiple phenotypes. CpGs with longitudinal mQTLs are more heritable and enriched in regulatory elements and pathways related to multicellular organism development and cell adhesion. These results chart dynamic genetic influences on the human methylome and provide a novel perspective on epigenetic regulation.

## Introduction

Extensive research has focused on the determinants of DNA methylation (DNAm), given its central role in development, ageing and human complex traits. It is now well established that DNAm changes across the life course^1^ and is influenced by both environmental and genetic factors^2,3^. Epigenome-wide association studies (EWAS) have revealed that more than half of the assayed CpG sites exhibit temporal changes throughout childhood, often following non-linear patterns^4^. Large-scale genome-wide association studies (GWAS) that treat methylation levels as quantitative outcomes aim to identify genetic loci underlying DNAm variation, termed methylation quantitative trait loci (mQTLs)^5^. Among the largest endeavours to date is the Genetics of DNA Methylation Consortium (GoDMC), which has established an extensive catalog of >270,000 independent mQTLs identified across 32,851 participants from 36 studies, assuming an additive genetic effect on DNAm^6^. Previous research based on DNAm measured at one time-point has suggested that age may modify genetic influences on DNAm, either through intrinsic aging processes or via interactions with environmental factors and variation in cellular proportions^7–9^. In addition, differences in mQTLs identified at different ages indicate the potential presence of time-varying associations^2^. However, such dynamic genetic effects on DNAm have not been systematically examined in a longitudinal context, largely due to the scarcity of datasets with repeated DNAm measurements from birth to adulthood.

In this study, we introduce the concept of longitudinal mQTLs, defined as single nucleotide polymorphisms (SNPs) whose influence on CpG methylation varies over time (Fig.1). Notably, this definition captures intra-individual variability in genetic effects, which is distinct from age interaction methylation QTLs (imeQTLs) that reflect inter-individual differences^8^. Additionally, CpGs with longitudinal mQTLs conceptually differ from epigenetic clock sites correlated with age^10^ or DNAm sites showing changes over time in longitudinal EWAS^4,11^, neither of which necessarily implicate underlying genetic variation. Gene regulation, characterized by its dynamic and adaptive nature, is considered central to adaptation, development, and health consequences, and age-dependent genetic influences on transcription have been implicated in age-related disorders^12,13^. Longitudinal birth cohort studies with repeated DNAm measurements on the same individuals and genetic data present a unique opportunity to examine the dynamic genetic effects on the epigenome, potentially yielding deeper mechanistic insights into the molecular pathways underlying human growth, development, and disease aetiology.

**Fig. 1.**
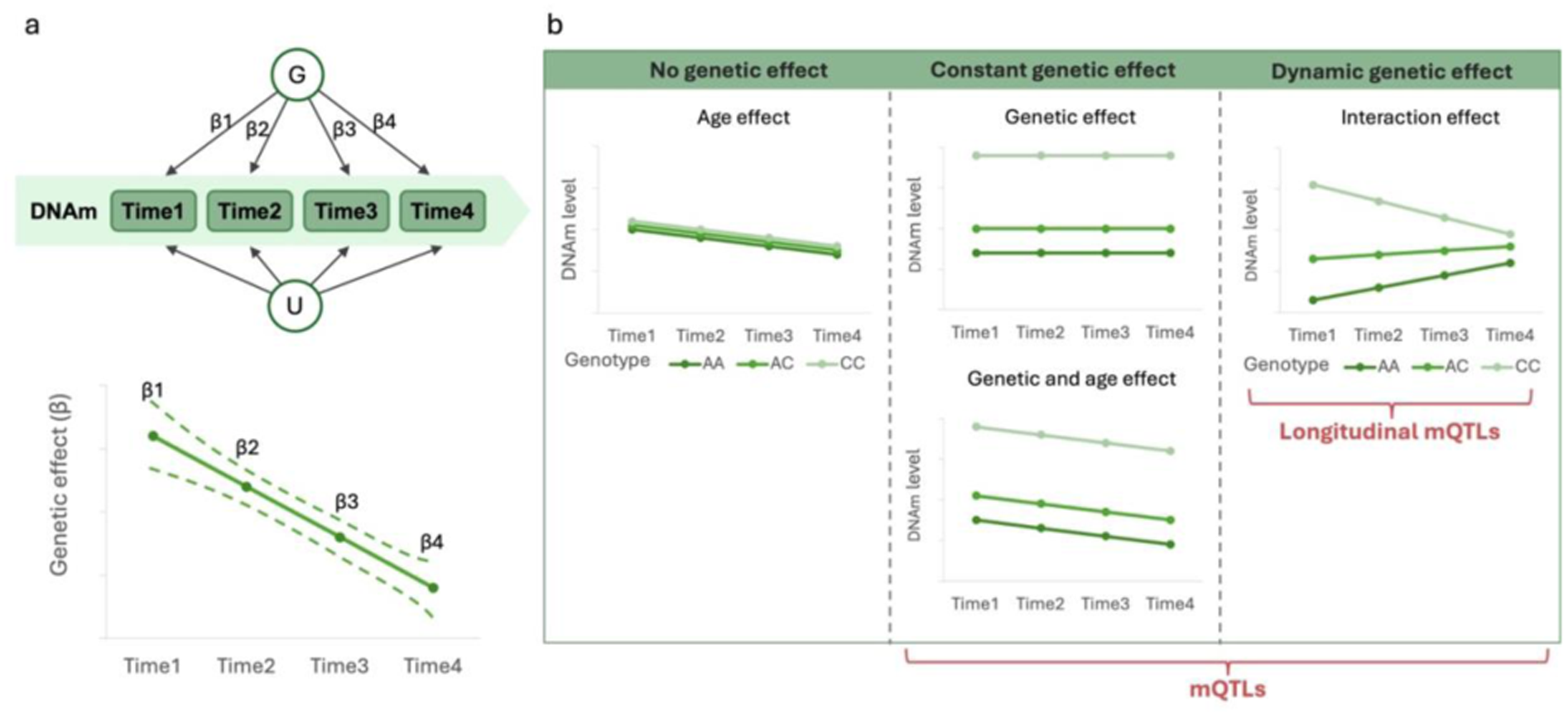
Graphical representation of longitudinal mQTLs. **a** Conceptual and statistical illustration of dynamic genetic influence on DNAm. G denotes genetic factors, and U represents potential confounders. **b** Trajectory plots differentiating CpG sites identified in studies focusing on age-related effects, genotype-related effects, and longitudinal mQTLs. mQTL = methylation quantitative trait locus.

This study aimed to identify genetic variants whose associations with blood DNAm change over time across three longitudinal cohort studies encompassing birth through early adulthood, and to explore the phenotypic implications and biological mechanisms underlying these dynamics.

## Results

### SNP-CpG pairs with dynamic associations

The discovery analysis used 3,416 blood samples collected from 1,057 participants of the Avon Longitudinal Study of Parents and Children (ALSPAC)^14^ who had repeated DNAm measurements spanning birth to early adulthood: birth (cord blood), and ages 7, 15, and 24 years (peripheral blood; Additional file 2: Table S1). Given the statistical power challenges inherent in detecting interaction effects^15^, we restricted our analysis to independent SNP-CpG pairs that were previously reported by GoDMC^6^ using life-course DNAm data as having additive mQTL associations (Fig.1; Additional file 1: Note S1). After applying quality control (QC) filters, 236,398 pairs were analysed, comprising 194,664 unique SNPs and 175,115 CpG sites (Additional file 1: Fig. S1). Of these, 20,141 pairs (8.5%) acted in *trans*.

We identified 2,393 longitudinal mQTL-CpG pairs with significant genotype-by-age interactions in linear mixed models (LMMs) at a Bonferroni-corrected threshold (*p* < 2 × 10^−7^; Additional file 2: Table S2). These signals involved 2,210 unique SNPs (1.14% of tested variants) and 2,329 unique CpGs (1.33% of tested sites). Fig. 2a shows the Manhattan plots and the genomic positional relationships between the identified SNPs and CpGs, while quantile-quantile (Q-Q) plot is shown in Additional file 1: Fig. S2. Among the significant associations, 2,217 longitudinal mQTL-CpG pairs were classified as *cis*-acting, and 176 (7.3%) as *trans*-acting.

**Fig. 2.**
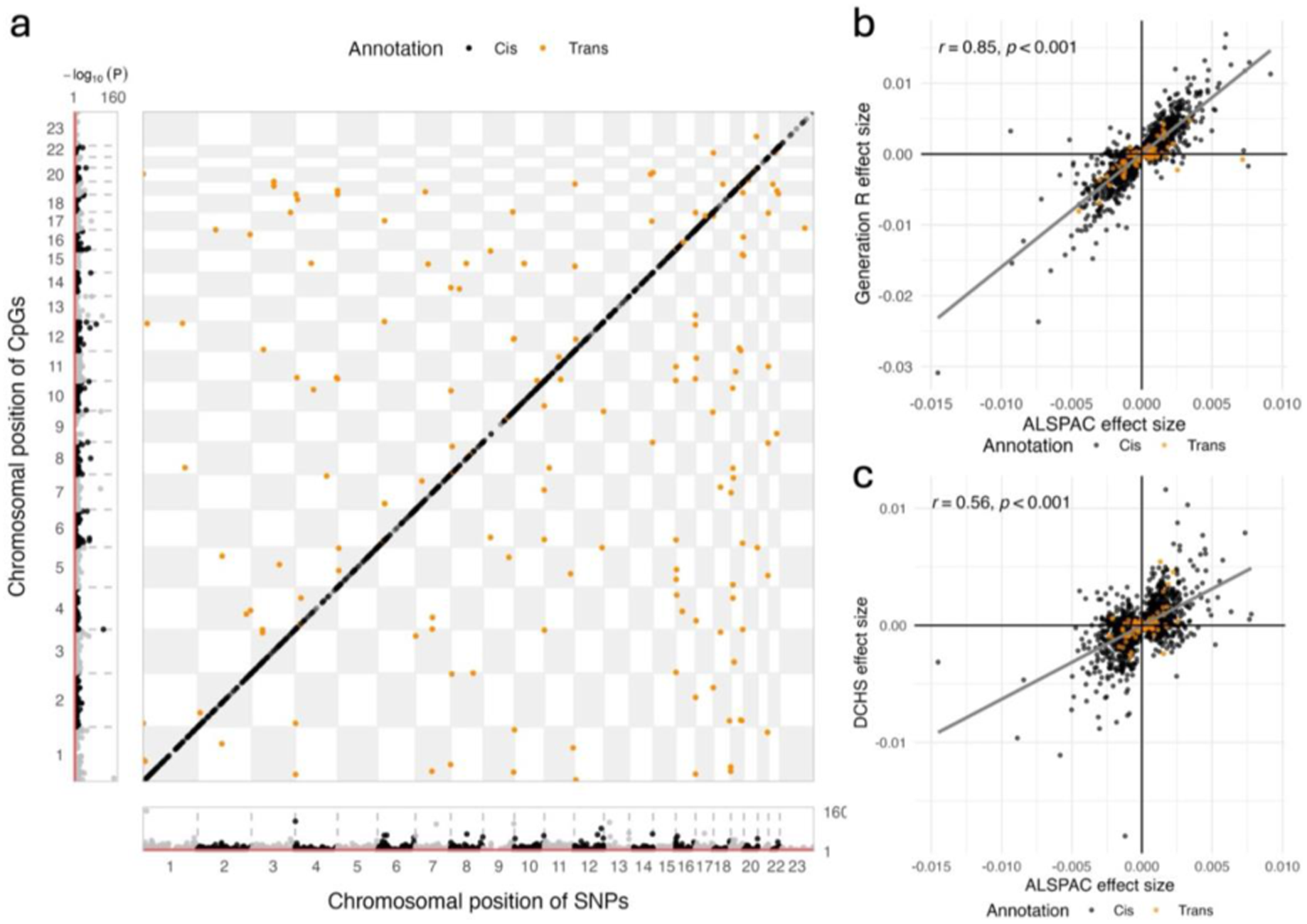
Discovery and replication of longitudinal mQTLs. Points are color-coded according to the *cis/trans* positional classification of mQTL-CpG pairs. **a** Chessboard plot showing the distribution of identified longitudinal mQTLs and their paired CpG sites. *Cis* mQTLs are indicated by black dots along the diagonal. Manhattan plots for tested SNPs and CpGs are shown at the bottom and left margins, respectively, where the red solid lines correspond to the significance threshold of *p* = 2 × 10^−7^. *P*-values are on a –log_10_ scale. **b** Effect size comparison of genotype-by-age interaction between the discovery cohort (ALSPAC) and the Generation R Study. The Pearson’s correlation coefficient (*r*) quantifies concordance across studies. **c** Effect size comparison of genotype-by-age interactions between the discovery cohort (ALSPAC) and the DCHS cohort. ALSPAC = Avon Longitudinal Study of Parents and Children, DCHS = Drakenstein Child Health Study.

We performed negative control and sensitivity analyses to assess the robustness of our methodology. As expected, no significant signals were detected among the 100,000 permutated SNP-CpG pairs with no prior evidence of association (Additional file 1: Fig. S3). Sensitivity analyses using complete cases, covariates excluding cellular composition, or samples from time points beyond infancy yielded results highly consistent with the main analysis (all Pearson’s *r* ≥ 0.69; Additional file 2: Table S3).

### Replication analysis

We conducted replication analysis of the 2,393 identified longitudinal mQTL-CpG pairs in two independent cohorts differing in age range and ancestry from ALSPAC — the Generation R Study and the Drakenstein Child Health Study (DCHS) — to assess the generalisability of our findings.

In Generation R, a total of 1,515 blood samples from 595 European children were included, with DNAm measured at a minimum of two time points: birth (cord blood), age 6, and age 10 (peripheral blood; Additional file 2: Table S1). Due to differences in genotyping chips across cohorts and QC procedures, only 1,862 SNP-CpG pairs were available for analysis, so a significance threshold of *p* < 2.7 × 10^−5^ was applied. The minor allele frequencies (MAFs) of these SNPs (Pearson’s *r* = 0.99) and their main effects on DNAm (*r* = 0.93) were highly consistent with those observed in ALSPAC (Additional file 1: Fig. S4). The estimated genotype-by-age interaction effects also correlated strongly with ALSPAC estimates (*r* = 0.85; Fig. 2b). 996 pairs were directionally consistent and Bonferroni-significant, exceeding the 441 pairs expected given power differences between cohorts^16^, indicating substantial reproducibility.

Replication in the DCHS cohort included 1,576 peripheral blood samples from 624 children of Black African and mixed ancestry, with DNAm measured at a minimum of two time points among ages 1, 3, and 5 (Additional file 2: Table S1). After QC, 1,798 SNP-CpG pairs remained, yielding a significance threshold of *p* < 2.8 × 10^−5^. Their SNP MAFs (Pearson’s *r* = 0.34) and main effects (*r* = 0.74) were less concordant with those in ALSPAC (Additional file 1: Fig. S4). Interaction effects showed a moderate correlation with the discovery cohort (*r* = 0.56; Fig. 2c). 146 pairs replicated with consistent signs and Bonferroni significance, compared with 190 expected after accounting for power differences.

Overall, longitudinal mQTL effects replicated reasonably well, with stronger out-of-sample consistency observed in Generation R, which had a greater overlap in age range and a closer ancestral match to the discovery cohort^17^. Interestingly, some SNP-CpG pairs showed stronger interaction effects in the replication cohorts than in ALSPAC. This is likely attributable to differences in age ranges across cohorts, as genetic effects sensitive to developmental timing may vary considerably at younger ages.

### Trajectories of identified dynamic associations

Amongst the 2,393 longitudinal mQTL-CpG associations, we identified four distinct trajectory patterns across the first 24 years of life, which reflected increasing, decreasing, fluctuating, or sign-reversed changes in the genetic effect on DNAm (Fig. 3; Additional file 1: Note S2). An increase in the absolute effect sizes was observed in 1,327 longitudinal mQTL-CpG pairs (55.5%; Additional file 2: Table S4). By contrast, 843 pairs (35.2%) showed decreasing effect sizes over time, while 100 (4.2%) exhibited fluctuating patterns, and 123 pairs (5.1%) underwent a switch in the sign of their β coefficients within the age range studied. The 0 to 7-year age span encompassed the most pronounced changes in the effects of longitudinal mQTLs (Additional file 1: Fig. S5). At birth, most SNP-CpG pairs demonstrated minimal associations. With increasing child age, a growing proportion of pairs exhibited stronger associations, indicating increasing genetic influence on DNAm during early development.

**Fig. 3.**
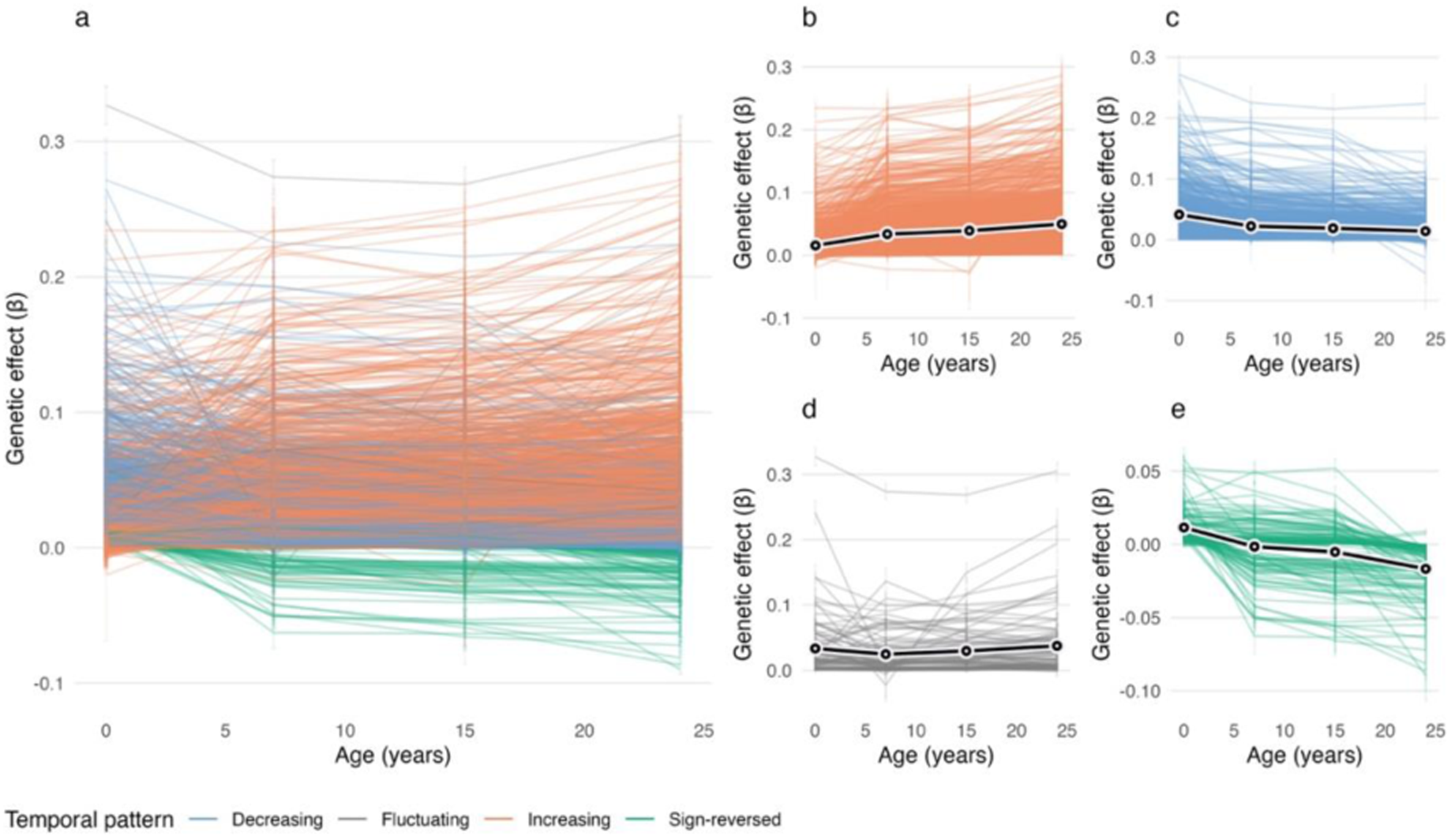
Trajectories of longitudinal mQTL-CpG associations grouped by temporal patterns. Each trajectory represents β coefficients (time-specific genetic effects on DNAm) with corresponding error bars at 4 time points. Curves are color-coded according to the temporal patterns of genetic influence. Black lines indicate the average genetic effects for each pattern. **a** Trajectories for all identified longitudinal mQTL-CpG pairs. **b** Trajectories with increasing absolute effect sizes over time. **c** Trajectories with decreasing absolute effect sizes over time. **d** Trajectories showing a fluctuating pattern. **e** Trajectories characterized by a reversal in the sign of genetic effects over time. Sign-reversed indicates positive genetic effects at some time points and negative effects at others.

These observations suggest that differences in sampling age ranges may partly account for the replication heterogeneity. For example, since DCHS samples were collected at ages 1–5, their estimates are likely to be more sensitive to genetic effect changes occurring in early life than those emerging after age 7. As expected, longitudinal mQTL-CpG pairs exhibiting a fluctuating pattern (Pearson’s *r* = 0.25) showed the poorest replication compared with other patterns (*r* ≥ 0.56; Additional file 2: Table S5).

### Phenotypic associations of longitudinal mQTLs

Comparing the MAF distribution of the 2,210 longitudinal mQTLs with the remaining 192,454 tested mQTLs revealed that low-frequency variants were underrepresented among longitudinal mQTLs and showed larger interaction effects (Additional file 1: Fig. S6). This pattern likely reflects the reduced statistical power to detect associations involving rare variants.

To assess the phenotypic relevance of the identified longitudinal mQTLs, we examined their reported associations with complex traits. Based on GWAS data accessed from the OpenGWAS database^18^, 835 longitudinal mQTL SNPs overlapped with SNPs associated with 589 UK Biobank phenotypes, spanning domains including physiological indexes, body composition, anthropometric measures, and health-related traits (Additional file 1: Fig. S7). This suggests that these SNPs may not only shape DNAm patterns but also contribute to adult phenotypes. Furthermore, we observed minimal overlap between our discovery longitudinal mQTLs and additive genetic associations for age-related traits, including childhood adiposity^19^, longitudinal growth traits^20^, and epigenetic age clocks^21^ (Additional file 2: Table S6). Similarly, overlap with variance QTL associations^22^ was negligible. Nevertheless, 55 longitudinal mQTLs overlapped with previously reported cell type imeQTLs^8^, a type of interaction QTL whose effects depend on leukocyte abundance, and 9 of them affected the same CpG site. Trajectories of the 9 longitudinal mQTL-CpG pairs revealed changing associations not only between ages 0 and 7 but also late in life, suggesting that this overlap was not solely attributable to differences in blood tissue types used (Additional file 1: Fig. S8). We also assessed overlap with development-related traits, such as age at menarche, early-life anthropometric traits, metabolic indicators, cell counts, hormonal factors, growth factors, and neurodevelopmental disorders (Additional file 2: Table S6). Notably, 31, 17, and 10 longitudinal mQTLs were linked to growth hormone receptor, BDNF/NT-3 growth factor receptor, and low-density lipoprotein (LDL) cholesterol, respectively, whereas overlaps with other traits were minimal (<10 mQTLs per trait).

We then performed enrichment analysis to assess whether the SNPs with dynamic genetic effects on DNAm contributed more significantly to development-related traits than other tested SNPs (additive mQTLs discovered in GoDMC). Despite their small magnitude, correlations between genotype-by-age effects on DNAm and genetic effects on heel bone mineral density (Pearson’s *r* = 0.0082) and glucose levels (*r* = 0.0076) were significant after Bonferroni correction across 31 traits (*p* < 1.6 × 10^−3^; Additional file 2: Table S7). The magnitude of SNP associations with LDL cholesterol or white blood cell counts (neutrophil, monocyte, basophil, eosinophil, and lymphocyte counts) was positively related to the likelihood of a SNP being longitudinal mQTLs (*p* < 1.6 × 10^−3^), whereas stronger genotype-by-age interaction effects on DNAm did not predict trait associations. Additionally, when comparing our results with two previous studies that investigated age-dependent genetic effects on complex traits and structural brain changes using cross-sectional data^23,24^, we found nominal correlations between genotype-by-age interaction effects on DNAm and those on BMI (*r* = -0.0047), weight (*r* = -0.0047), cerebral white matter (*r* = 0.0055), and pallidum (*r* = -0.0051; *p* < 0.05; Additional file 2: Table S8).

Overall, despite substantial overlap with GWAS signals across a variety of traits, identified longitudinal mQTLs showed more pronounced enrichment for only a few traits compared to GoDMC mQTLs, which may themselves include undetected longitudinal effects due to limited power.

### CpGs with longitudinal mQTLs

Compared to the other 172,786 tested CpGs, the 2,329 longitudinal mQTL CpGs were characterized by lower DNAm levels and greater variability (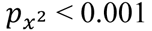; Additional file 1: Fig. S9). Among these, 1,077 (45.0%) CpGs had low methylation levels (β value: 0-0.2), and 412 (17.2%) had high methylation levels (β value: 0.8-1).

Next, we analyzed the variance composition of longitudinal mQTL CpGs using variance decomposition estimates derived from a twin-based study of individuals aged 17–79 years^9^. CpGs with longitudinal mQTL associations exhibited significantly higher heritability compared to other tested sites (mean [standard deviation] of total additive heritability: 0.58 [0.26] vs. 0.36 [0.21]; t-test: *p* < 0.001) (Fig. 4). Moreover, stratifying CpGs by the trajectory patterns of their genetic effect revealed notable differences (𝑝_ANOVA_ < 0.001): those with increasing genetic influence over time displayed the highest mean heritability (0.66 [0.24]), while CpGs with fluctuating effects had the lowest (0.47 [0.29]).

**Fig. 4.**
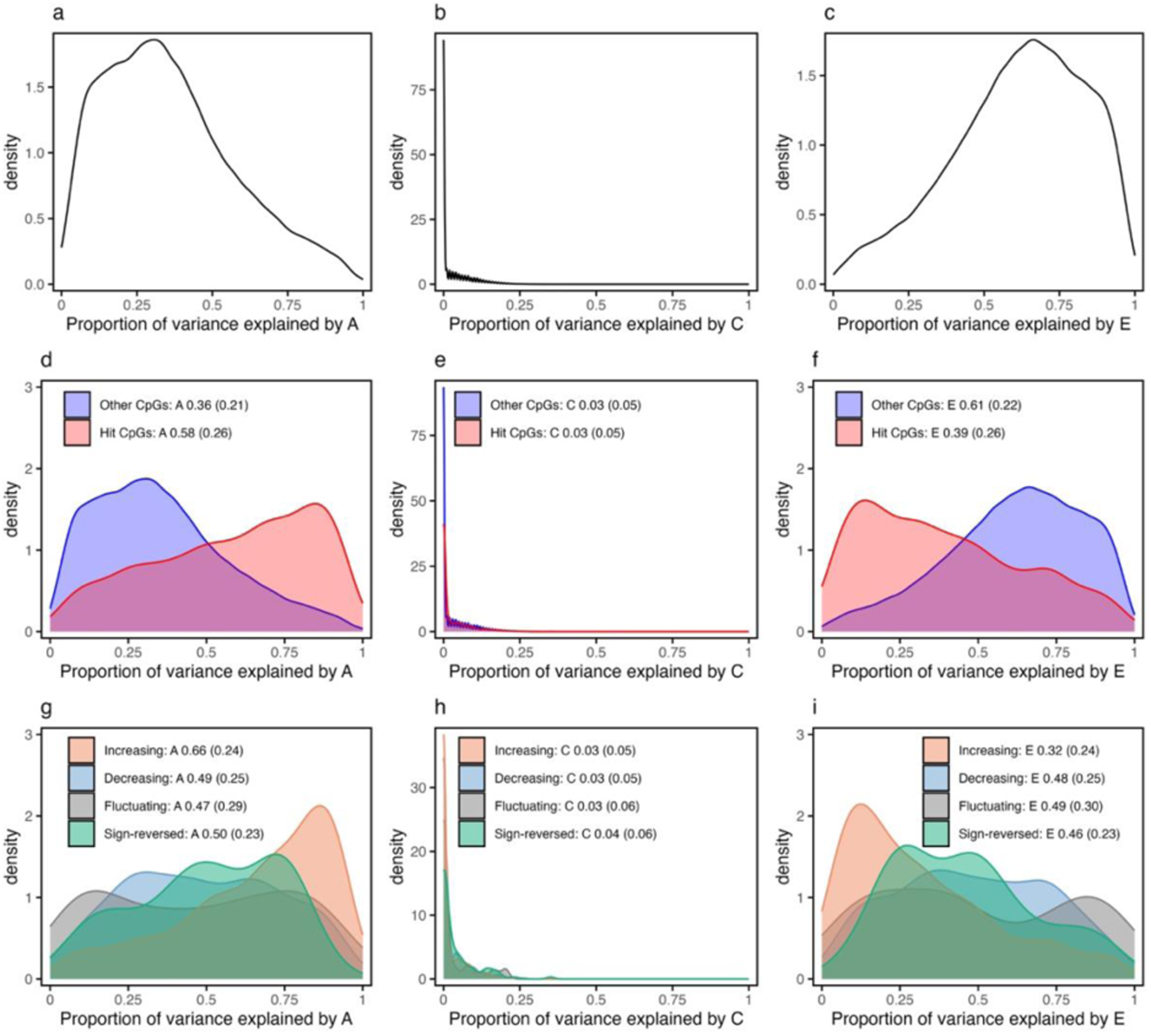
Comparison of variance decomposition estimates between CpGs with mQTLs and those with longitudinal mQTLs. Numbers in the legends represent the mean (SD) proportion of the variance in DNAm explained by additive genetic effects (A), common environmental effects (C) and unique environmental factors including measurement error and stochastic variation (E), across different sets of CpG sites. Information for 41,141 tested CpGs, including 440 identified CpGs, was missing due to data unavailability or unreliable heritability estimates. **a-c** Variance estimates for 133,974 tested CpGs, each associated with at least one SNP in the GoDMC database. **d-f** Variance estimates of tested CpGs, grouped by whether the CpGs exhibited a longitudinal association with at least one SNP (hit CpGs) or did not (other CpGs). **g-i** Variance estimates for 1,909 longitudinal mQTL CpGs grouped by temporal trajectory patterns. SD = standard deviation, GoDMC = Genetics of DNA Methylation Consortium.

Some overlap was observed between the identified CpGs and sites reported in previous DNAm studies (Additional file 2: Table S9). Of the 2,329 CpGs associated with longitudinal mQTLs, 1,733 CpGs demonstrated linear or non-linear associations with chronological age in a pediatric age EWAS (with some of the sample drawn from ALSPAC, 289,051 hit sites)^4^, while only 25 showed age-related changes in an adult EWAS (≥49 years, 1,316 hit sites)^25^. As Fig.1b shows, CpGs with dynamic genetic associations do not always exhibit clear population-level changes over time, as these effects may be offset by opposing genotype-specific trends, confined to certain age ranges, or driven by prevalent genotypes (Additional file 1: Fig. S10). Epigenetic clock studies, which aim to identify DNAm sites predictive of chronological or biological age across individuals irrespective of genetic variation, also fundamentally differ from this research, which investigates the genetic determinants of intra-individual DNAm changes over time and development. Indeed, only 1, 1, 9, and 2 of our identified CpG sites were included in Horvath’s clock^26^, Hannum’s clock^27^, PhenoAge^28^, and DunedinPACE^29^ respectively. Additionally, 428 of the 2,329 identified CpGs overlapped with previously reported cell type imeQTL CpGs, and 1 overlapped with an age imeQTL CpG site^8^. According to exposure EWASs, 81 longitudinal mQTL CpGs were linked to maternal smoking during pregnancy^30^, of which 44 and 34 displayed decreasing and increasing postnatal genetic influences, respectively.

### Functional enrichment

CpGs associated with longitudinal mQTLs were enriched in CpG islands and depleted in open sea regions, relevant to the other CpGs (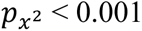; Additional file 1: Fig. S8). In addition, they were preferentially located within regulatory elements across a wide range of tissues and cell types (Additional file 1: Fig. S11). Specifically, they were strongly overrepresented in DNase I hypersensitive sites across nearly all examined cell types, as well as H3K4me1, H3K4me3, and H3K27me3 histone modifications. They were also more likely to localize to promoter, enhancer, and repressed PolyComb regions^31^. This enrichment was even more pronounced compared to that observed for equivalent numbers of genotype-associated or age-associated CpGs (Additional file 1: Fig. S12 and Fig. S13).

We also conducted functional enrichment analysis at gene level. Of the identified CpGs, 1,690 were successfully mapped to 1,423 genes. These genes exhibited enrichment in regulatory motifs of multiple transcriptional factors (TFs) and biological pathways broadly related to development (anatomical, nervous, circulatory, and muscle systems), homeostasis and metabolism, cellular localization and transport, and regulation of insulin secretion (Fig. 5a; Additional file 2: Table S10). Comparison with 18,664 genes linked to GoDMC mQTL CpGs still revealed overrepresentation in regulatory motifs and pathways associated with multicellular organism development and cell adhesion (Fig. 5b; Additional file 2: Table S11).

**Fig. 5.**
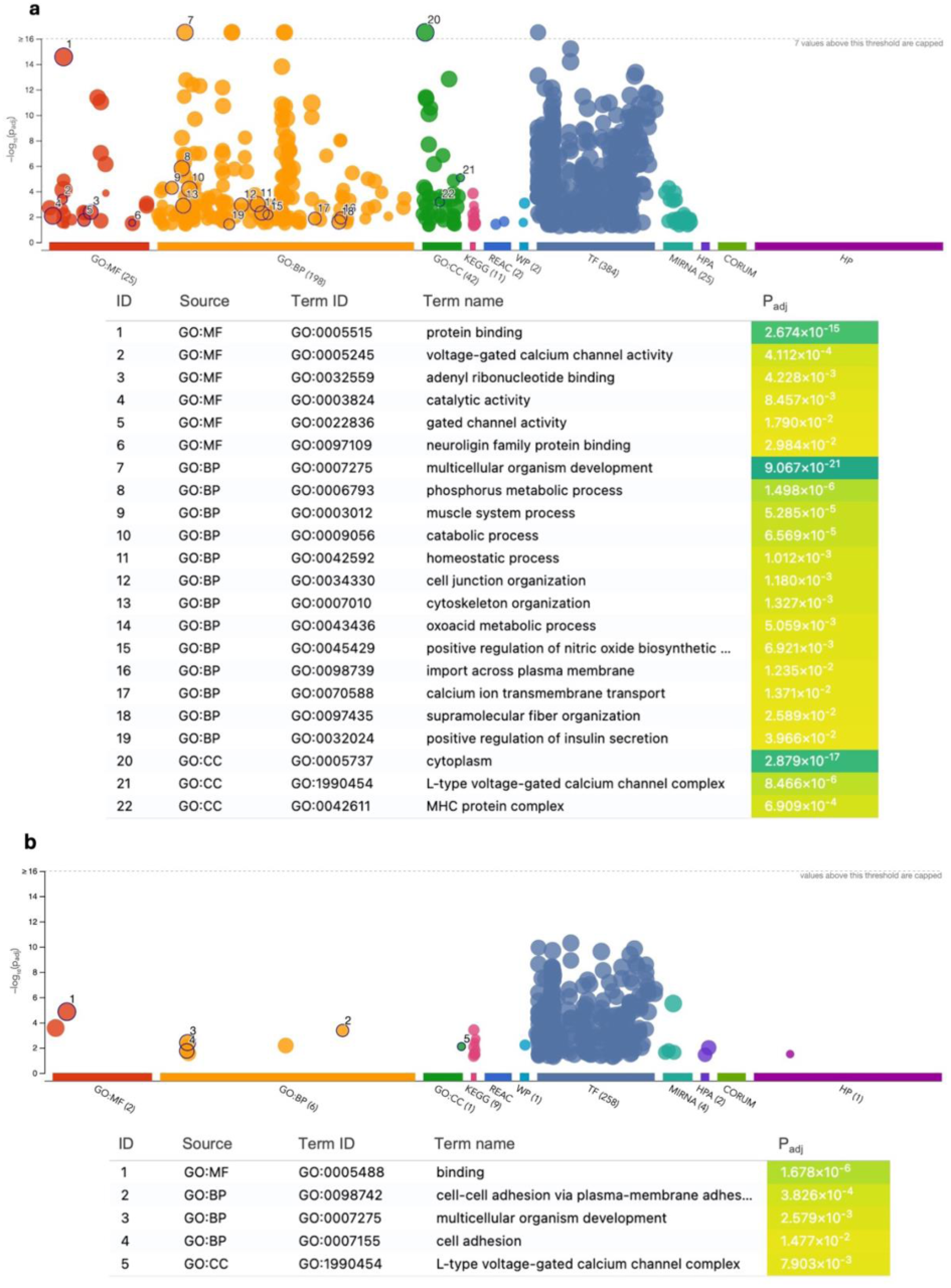
Functional enrichment of 1,423 genes mapped to longitudinal mQTL CpGs. The x-axis denotes different data sources: Gene Ontology (GO), KEGG, Reactome (REAC), WikiPathways (WP), TRANSFAC (TF), miRTarBase (MIRNA), Human Protein Atlas (HPA), CORUM, and Human Phenotype Ontology (HP). The y-axis represents the –log₁₀ transformed *p*-values, which were adjusted for multiple testing using the g:SCS algorithm at a significance threshold of 0.05. Highlighted dots for GO enrichment results represent leading terms, core enriched terms likely driving the significance of nearby related terms, thereby enhancing interpretability while considering the hierarchical structure of the GO. **a** Enrichment analysis using all annotated genes as the background. **b** Enrichment analysis using genes mapped to GoDMC mQTL CpGs as the background. MF = molecular function, BP = biological process, CC = cellular component.

While GoDMC reported an overrepresentation of TF-binding sites for CTCF, SMC3, and RAD21 among *trans*-mQTL CpGs^6^, our analysis identified enrichment exclusively for CTCF, a key architectural protein involved in 3D genome organization and the regulation of enhancer–promoter interactions^32,33^.

## Discussion

This study systematically investigated genotype-by-age interactions on DNAm during the first two decades of life. We identified a limited number of longitudinal mQTLs (1.14% of ∼200,000 tested variants), whose effects on DNAm varied significantly over time. Despite the modest proportion, these mQTLs exhibited consistent longitudinal effects across cohorts of different ancestries and age ranges. Moreover, the longitudinal mQTLs were categorised in four temporal trajectories, where the trajectory with increasing absolute genetic effects with age was the most prevalent pattern. Changes in mQTL effect size were most pronounced during the early-life period (0–7 years). Compared to those exhibiting stable effects, SNPs and CpGs involved in longitudinal associations displayed distinct molecular and functional features. Longitudinal mQTLs overlapped with SNPs associated with diverse human traits, showing notable relevance to heel bone mineral density, glucose levels, LDL cholesterol, and white blood cell counts. CpGs with longitudinal mQTLs showed a greater contribution of genetic factors to methylation variance than did other mQTL CpGs. Furthermore, they were significantly enriched in gene expression regulatory elements as well as biological pathways related to organismal development and cell adhesion. Together, these findings suggest that genotype-by-age interactions, although uncommon, may underlie developmental processes.

Our results align with previous studies showing that genetic control of DNAm remains mainly stable over time. A previous mQTL study found that around 95% of mQTLs identified at a specific age also presented detectable effects at other ages, while a smaller proportion (85%) of mQTLs discovered at birth were successfully replicated in later life^2^. A family-based study^34^ revealed age moderated genetic effects at 2.1% (7,806) of DNAm sites in 2,603 twin individuals, particularly in CpG islands and hypomethylated regions^9^, consistent with our observations. In addition, early life is a period critical for cellular differentiation and immune maturation. Prior genome-wide DNAm analyses have shown that numerous DNAm sites experience substantial reprogramming after birth before stabilizing later in life^1,4,35^. Together, the current evidence supports the robustness of cross-sectional mQTL discovery yet also emphasizes the need for developmental-stage-specific mapping, particularly in infancy. Notably, some stable mQTLs may still interact additively with age, producing compound effects over time that are not captured by our interaction models (Fig. 1b).

Genotype-by-age interaction effects on DNAm showed strong replication in the European cohort for the identified longitudinal mQTLs. The comparatively lower replication in the non-European cohort may partly reflect its narrower age range and allele-frequency differences, rather than ancestry per se. Previous studies indicate that most *cis*-mQTL effect sizes and methylation risk scores are broadly shared across ancestries, although some ancestry-specific effects exist^36,37^. These replication findings underscore the potential to improve power to detect dynamic genetic effects by meta-analysing populations sampled within comparable age ranges.

The temporal patterns observed in longitudinal mQTLs are likely driven by three plausible mechanisms that may act simultaneously and in concert. First, some longitudinal mQTLs may causally influence DNAm in an age-dependent manner (genome → methylome). This mechanism is supported by the increasing associations observed in half of the longitudinal mQTL-CpG pairs, as well as by the skewed distribution of identified CpGs toward highly heritable sites. Such time-varying influences may arise from gradual amplification of genetic effects over time^38^ or from normal developmental processes, including critical periods when specific genes exert differential effects^39,40^. Second, longitudinal mQTL-associated CpGs may play active roles in coordinated regulatory networks (methylome ↔ gene expression). They and their linked genes appear to be not only co-regulated by upstream transcriptional regulators but also involved in regulating downstream physiological and biochemical processes, forming multi-layered regulatory architectures that may underlie the observed dynamic genetic associations. Third, genotype-by-age effects may arise from differential environmental responsiveness (environment → methylome). Some of our identified CpGs were also influenced by maternal smoking during pregnancy, and individuals with different genotypes may respond uniquely to shared exposures^41,42^.

This study has several strengths and limitations. To our knowledge, this is the first study to directly profile age-dependent genetic effects on the epigenome. The availability of repeated DNAm measurements at multiple time points enabled a high-resolution assessment. By focusing on known mQTLs, we reduced the multiple-testing burden and partially alleviated the power constraints inherent in interaction studies^15^. Good replication in independent cohorts of diverse ancestries, together with successful negative control analyses, lends further credibility to our findings. We also acknowledge three main limitations. First, additional longitudinal mQTLs could be identified with larger sample sizes or a genome-wide search, especially those acting in *trans*. Our analysis was restricted to previously reported mQTL-CpG pairs for computational feasibility, but some heritable CpGs may have been missed due to reliance on additive models or because their genetic associations occur only during specific periods. Second, cord-peripheral blood differences may remain a potential source of bias, although we accounted for leukocyte heterogeneity. While the observed drift in genetic effects likely reflect biological processes intrinsic to post-birth development, tissue composition cannot be entirely ruled out. A unified reference panel for estimating leukocyte fractions across the life course became available during the preparation of this manuscript and may be valuable for future longitudinal DNAm studies^43^. Nevertheless, results from negative controls, supplementary analyses, and replication in peripheral blood indicate that such tissue differences are unlikely to meaningfully impact our conclusions. Third, our reliance on linear additive models does not accommodate complex forms of genetic regulation (dominance effects, epistasis, or parent-of-origin effects) or non-linear DNAm changes^5,44^.

Our findings raise several questions for future research. Given the role of DNAm as a mediator of genetic risk for complex traits, it is pertinent to ask to what extent the genotype-specific patterns of DNAm change can predict corresponding differences in phenotypic development or disease risk^38^. In addition, it would be informative to assess whether parental genotypes have an intergenerational impact on children’s DNAm dynamics^45,46^. Furthermore, studies incorporating repeated DNAm measurements across adult life stages and in other tissues are needed to assess the generalizability of these findings in older populations and beyond peripheral blood.

In conclusion, our study provides direct evidence for genotype-by-age interactions on DNAm from birth to adulthood. Although such dynamic genetic effects were observed at a limited number of loci, they replicated well across independent cohorts and exhibited phenotypic and development relevance. The trajectories that these dynamic effects follow are likely shaped by a combination of intrinsic genetic programs, regulatory complexity, and gene-environment interplay.

## Methods

### Longitudinal mQTL discovery study participants

This study used data from child participants in ALSPAC. This cohort recruited pregnant women living in Avon, United Kingdom with expected dates of delivery between 1991 and 1992 and collected blood samples from a subset of the mother and child pairs^47^. A total of 13,988 children from the initially enrolled pregnancies were alive at one year of age.

Additional pregnancies were recruited when the oldest children reached approximately seven years, resulting in a total of 15,447 pregnancies, of which 14,901 children were alive at one year of age. Its design and implementation have been comprehensively documented elsewhere^14,48–50^. The study website contains details of all the data that is available through a searchable data dictionary and variable search tool^51^. Only children were included in the analysis, as DNAm change patterns may differ substantially between mothers and children due to their distinct age groups. Participants of non-European ancestry or with DNAm data available at fewer than two time points were excluded from the analysis.

### Genetic data and epigenetic data

For ALSPAC child participants, genome-wide genotype data were imputed to GRCh37, and DNAm data were obtained from cord blood at birth and peripheral blood at age 7, 15 and 24 (Additional file 1: Note S1). After rigorous QC, approximately 7 × 10^6^ genetic variants and 4 × 10^5^ CpG probes remained. Rather than interrogating trillions of potential SNP-CpG permutations (7 × 10^6^ variants × 4 × 10^5^ probes), we focused on 248,607 *cis*-mQTLs and 23,117 *trans*-mQTLs that displayed significant associations with their corresponding CpGs in GoDMC^6^, ensuring analytical feasibility and sustainability (Additional file 1: Fig. S1). GoDMC represents the largest GWAS of genetic determinants of CpG methylation on the Illumina Infinium HumanMethylation450 BeadChip (HumanMethylation450 array), using DNAm data collected from birth through adulthood.

### Covariates

We included age, sex, methylation batch, the first ten genetic principal components (PCs), and estimated cellular composition as covariates. Age was calculated from participants’ birth dates and sample collection dates. Cellular composition was estimated at each time point through deconvolution of 12 leukocyte subtypes using the Salas reference panel^52^, including monocytes, neutrophils, eosinophils, basophils, natural killer cells, naïve and memory B-lymphocytes, naïve and memory CD4 + and CD8 + T-lymphocytes, and T regulatory cells. Of these, 11 were included as covariates. Although reference panels specifically designed for umbilical cord blood^53,54^ were available, we applied the Salas method across all samples to ensure consistency in cell type estimation across time points.

### Statistical analysis

The main analysis used two models. First, a linear model was fitted to 236,398 previously reported mQTL-CpG pairs, pooling data across all time points to estimate SNP main effects on DNAm. Secondly, a LMM was applied to evaluate the interaction between SNP and age on DNAm levels, using the lmerTest R package. The model was formulated as follows:

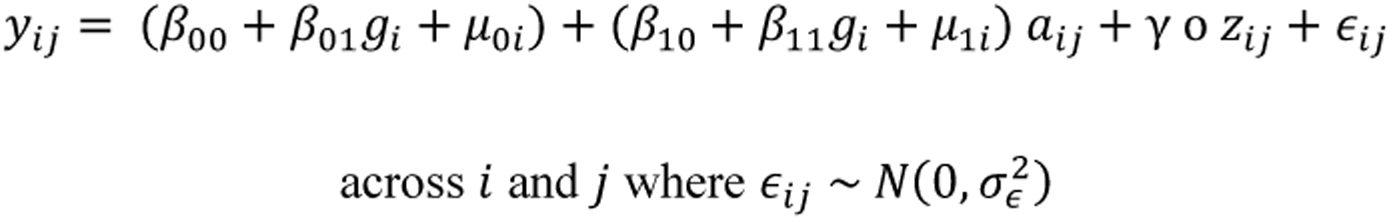

In this equation, 𝑦_𝑖𝑗_ denotes the DNAm level for individual 𝑖 at time point 𝑗 for a given CpG site; 𝑔_𝑖_ is the genotype of individual 𝑖 at the corresponding SNP; 𝑎_𝑖𝑗_ is the age of individual *i* at time point *j*; 𝛾 о 𝑧_𝑖𝑗_ denotes the Hadamard product between covariate effects and the covariate vector; 𝜇_0𝑖_ represents a random intercept accounting for inter-individual differences in DNAm levels at birth; 𝜇_1𝑖_ specifies a random slope for age; 𝜖_𝑖𝑗_ represents the residual errors. The parameter 𝛽^_11_ captures the longitudinal genotype-by-age interaction effect.

Both the linear model and the LMM were adjusted for the covariates stated above. SNPs were classified as longitudinal mQTLs if they reached significance in both models to reduce the likelihood of spurious interactions arising from genotype imbalance or non-linear DNAm trajectories. The significance threshold was adjusted for the number of tested SNP-CpG pairs using Bonferroni correction to *p* < 2 × 10^−7^. Longitudinal mQTLs were considered to act in *cis* if the corresponding CpGs were located within 1 Mb of the SNP on the same chromosome.

A negative control analysis was conducted to assess the potential for spurious associations arising from collider bias or variation in blood sources. Specifically, 100,000 mQTL-CpG pairs from the analysis set were randomly selected and shuffled, creating a synthetic dataset comprising SNPs and CpGs with no known relationship. The same LMM was then applied to this randomized set.

Sensitivity analyses included: (i) exclusion of participants who did not have DNAm data at all four time points, (ii) analyses without adjustment for cellular composition, and (iii) exclusion of DNAm data at birth, which were derived from cord blood samples.

All analyses were primarily carried out in Python (version 3.10) and R (version 4.3.0). Unless otherwise specified, a *p*-value threshold of 0.05 was applied.

### Replication in independent birth cohorts

To validate the significant longitudinal mQTL associations identified in this study, replication was performed using data from two independent birth cohorts: the Generation R Study and the DCHS. The Generation R Study enrolled pregnant women with a delivery date between April 2002 and January 2006 in the study area of Rotterdam, Netherlands^55^. Blood samples from child participants were collected at birth, age 6, and age 10. Like in ALSPAC, their DNAm measurements at birth came from cord blood and at later time points from peripheral blood. Therefore, we sought birth cohorts with peripheral blood samples collected during early life to fill the gap between ages 0 and 5. The DCHS recruited pregnant women in Paarl, South Africa^56^. Peripheral blood samples from children were available at ages 1, 3, and 5, enabling examination of the dynamic genetic effects in the same blood tissue. Nevertheless, DCHS participants were ethnically and culturally heterogeneous, comprising approximately equal proportions of individuals of Black African and mixed ancestry.

For Generation R, replication analyses followed the same QC procedures and modelling approach as the discovery analysis. For DCHS, replication analyses were additionally adjusted for clinic site and ethnicity to account for study-specific sources of variation, and leukocyte subtypes were estimated using the Houseman method^7^. *P*-value thresholds were defined as 0.05 divided by the number of available SNP-CpG pairs in each cohort (Additional file 1: Note S1). Replication was assessed based on the Pearson’s correlation (*r*) between estimated interaction effects (genotype × age) in the discovery and replication datasets, as well as by comparing observed replication results with expected values. Expected values were the anticipated numbers of SNP-CpG pairs showing consistent effect directions or recurrent significance, accounting for differences in statistical power between cohorts using their standard error ^16^.

### Trajectories of genetic influences on DNAm

For the identified longitudinal mQTLs, time-specific genetic effects on DNAm were estimated at each time point using a linear model adjusted for the same covariates as in the main analysis. To facilitate comparison, trajectories with negative initial values were flipped, so that all trajectories had positive initial values. Trajectories were then classified into four categories based on the temporal patterns of genetic influence (Additional file 1: Note S2). Sankey plots were employed to exhibit the extent of transitions in genetic effects on DNAm from birth through to adulthood.

### Enrichment of longitudinal mQTLs

To investigate the phenotypic relevance of the longitudinal mQTLs identified in this study, we queried their association with UK Biobank phenotypes from the OpenGWAS database^18^. We also compared our results with genetic association studies of age-related and development-related traits, including anthropometric, biochemical, physiological, haematological, health, and brain imaging measures^8,18–24^. For traits without full GWAS summary statistics, we examined whether their associated loci overlapped with, or were in high linkage disequilibrium (LD; r^2^ > 0.8) with, longitudinal mQTLs. For traits with publicly available GWAS summary statistics, we computed Pearson’s correlations between effect estimates and applied a binomial regression model to assess enrichment (Additional file 1: Note S3). Significance was evaluated at *p* < 1.6 × 10^−3^ after Bonferroni correction.

### Decomposition analysis of DNAm variance

To examine the extent of genetic contribution to DNAm variation across different CpG site categories, we downloaded total heritability estimates and twin correlation for DNAm sites in blood from a previously published study employing the classical ACE twin model ^9^. This study analyzed 411,169 DNAm sites in peripheral blood from 769 monozygotic (MZ) and 424 dizygotic (DZ) twin pairs, aged 17–79 years, from the Netherlands Twin Register biobank. CpG sites with biologically implausible or statistically unreliable heritability metrics (e.g., heritability estimates > 1, negative intra-pair correlation coefficients) were discarded. We used t-tests and ANOVA to compare heritability estimates across different CpG subgroups.

### Functional enrichment analysis

To assess the functional relevance of the identified CpGs, we evaluated whether they were enriched in key chromatin-based epigenetic features that regulate gene expression. Specifically, we evaluated their overlap with DNase I hypersensitive sites, histone modification marks, and chromatin states across a broad range of cell types using eFORGE v2.0^57^. These regulatory elements are indicative of chromatin accessibility, gene regulatory activity, and transcriptional potential, respectively. To provide a reference for comparison between CpGs with longitudinal mQTLs and other types of DNAm sites, we applied the same enrichment pipeline to an equal number of randomly selected mQTL CpGs from GoDMC ^6^ and time-varying CpGs from a longitudinal EWAS^4^.

Gene functional profiling was performed using g:Profiler (version: e113_eg59_p19_f6a03c19)^58^. CpG-to-gene mapping was based on probe locations from the HumanMethylation450 array annotation. Enrichment analysis was performed across multiple databases, including Gene Ontology (GO: molecular function, biological process, and cellular component), pathway annotations (KEGG, Reactome, and WikiPathways), regulatory motifs in DNA (TRANSFAC), miRNA targets (miRTarBase), protein datasets (Human Protein Atlas and CORUM), and the Human Phenotype Ontology. The statistical domain scope was set to only annotated genes or gene lists identified for GoDMC hits.

## Declarations

### Ethics approval and consent to participate

Ethical approval for the study was obtained from the ALSPAC Ethics and Law Committee and the Local Research Ethics Committees. Written informed consent has been obtained from all ALSPAC participants. Consent for biological samples has been collected in accordance with the Human Tissue Act (2004). Study participation is voluntary and during all data collection sweeps, information was provided on the intended use of data. The Generation R Study is conducted in accordance with the World Medical Association Declaration of Helsinki and has been approved by the Medical Ethics Committee of Erasmus MC, Rotterdam. Informed consent was obtained for all participants. The DCHS was approved by the faculty of Health Sciences, Human Research Ethics Committee, University of Cape Town, Stellenbosch University and the Western Cape Provincial Health Research committee. Expecting mothers/guardian provided informed written consent at enrolment and were reconsented annually following childbirth.

### Availability of data and materials

The quality control procedures and analysis pipeline were shared through GitHub (https://github.com/MRCIEU/longitudinal_mQTL).

All ALSPAC cohort data used in this study were provided to the authors via their standard Access Policy. The informed consent obtained from ALSPAC (Avon Longitudinal Study of Parents and Children) participants does not allow the data to be made available through any third party maintained public repository. Supporting data are available from ALSPAC on request under the approved proposal number, B4401. Data are available to other bona fide researchers on the same basis; full instructions for applying for data access can be found here: http://www.bristol.ac.uk/alspac/researchers/access/. The ALSPAC study website contains details of all available data (http://www.bristol.ac.uk/alspac/researchers/our-data/). Data from Generation R are available upon reasonable request to its director, subject to local, national and European rules and regulations. Researchers who are interested in DCHS datasets can contact the PIs, Heather Zar and Dan Stein with a concept note outlining the request. More information can be found on their website (http://www.paediatrics.uct.ac.za/scah/dclhs).

## Competing interests

None. JRS is currently employed by UCB Pharma SA, but his contribution to this research was made during his time at the University of Bristol, prior to joining the company.

## Acknowledgements

This research was funded in whole by the Wellcome Trust (Grant ref: 317540/Z/24/Z) and carried out using the High Performance Computing facility and the Research Data Storage Facility of the Advanced Computing Research Centre, University of Bristol (http://www.bris.ac.uk/acrc/).

We are extremely grateful to all participants involved in this study, to the midwives for their help in recruiting them, and to the dedicated teams behind ALSPAC, Generation R, and the DCHS cohorts, which includes data collection staff, data and administrations staff, technical managers and the technical staff. We also would like to thank Richard Parker and Helen Natukunda for their assistance in examining genetic loci overlap with longitudinal traits.

JM is supported by MR/X021599/1. GH, KT, VW and TRG were in part supported by the Integrative Epidemiology Unit, which receives funding from the Medical Research Council and the University of Bristol (MC_UU_00032/1, MC_UU_00032/2, MC_UU_00032/3). GH, TRG and KT were supported by the National Institute for Health and Care Research (NIHR) Bristol Biomedical Research Centre (NIHR 203315). JRS work was supported by an MRC Methodology Research Grant (MR/M025020/1). AG is funded by Research Ireland through the Research Ireland Centre for Research Training in Genomics Data Science (18/CRT/6214) and by the EU’s Horizon 2020 research and innovation programme under the Marie Sklodowska-Curie grant H2020-MSCA-COFUND-2019-945385. AAL is supported by an MQ Fellows Award (MQF22\9). The work of CAMC is supported by the European Union’s Horizon Europe Programme (FAMILY, grant agreement No 101057529; HappyMums, grant agreement No 101057390) and the European Research Council (TEMPO; grant agreement No 101039672). This research was conducted while CAMC was a Hevolution/AFAR New Investigator Awardee in Aging Biology and Geroscience Research. DJS and HJZ are funded by the SAMRC.

## Authors’ contributions

YL: Software, formal analysis, data curation, writing – original draft, writing – review & editing, visualization, project administration. JRS and KT: Conceptualization, methodology. AG and CC: Data curation, formal analysis, validation, writing – review & editing. AAL, JFF, CAMC, DJS, HJZ, RM, and AJS: Resources, supervision, writing – review & editing. ECD, VW and TRG: Supervision, funding acquisition. GH and JLM: Conceptualization, methodology, software, data curation, resources, writing – review & editing, supervision, project administration. All authors read and approved the final manuscript.

