## Supplemental Notes and Figures for "Identification of Dynamic Genetic Influences on DNA Methylation from Birth to Adulthood"

This supplementary document accompanies the manuscript titled “*Identification of Dynamic Genetic Influences on DNA Methylation from Birth to Adulthood*”

### Note S1. Genetic and DNA methylation data in discovery and replication cohorts

**Avon Longitudinal Study of Parents and Children (ALSPAC)**

ALSPAC participants were genotyped based on the Illumina HumanHap550 BeadChip. The genotype data were imputed to the Haplotype Reference Consortium panel (hg19/GRCh37 human reference genome) and underwent rigorous quality control in accordance with standard best practices (1). SNPs with a minor allele frequency (MAF) > 0.01 and imputation quality > 0.8 were retained.

Genome-wide DNAm data were generated using the Illumina Infinium HumanMethylation450 BeadChip (450k array) for blood samples collected at birth, age 7, and age 15, and using the Illumina MethylationEPIC BeadChip (EPIC array) for samples collected at age 24. Probes that mapped to multiple genomic loci were excluded from analyses. CpG sites that are present only on the EPIC array but not on the 450k array were also eliminated, as these sites lack repeated measures across the longitudinal time points. Functional normalization was applied to the repeated methylation measures to minimize technical variation, and the measures were scaled to β-values ranging from 0 (unmethylated) to 1 (fully methylated). β-values exceeding three SDs from the time-point specific mean were identified as outliers and removed. Probes were annotated to gene-centric regions and regions related to CpG islands — CpG island, N shelf/shore, S shore/shelf, and open sea — based on the Illumina 450k array v1.2 manifest (2).

The UK Medical Research Council and Wellcome (Grant ref: MR/Z505924/1) and the University of Bristol provide core support for ALSPAC. A comprehensive list of grants funding is available on the ALSPAC website (http://www.bristol.ac.uk/alspac/external/documents/grant-acknowledgements.pdf); Data used in this research was specifically funded by the Biotechnology and Biological Sciences Research Council (BBSRC, Grant ref: BBI025751/1 and BB/I025263/1), the MRC Integrative Epidemiology Unit (IEU, Grant ref: MC_UU_12013/1 & MC_UU_12013/2 & MC_UU_12013/8), National Institute of Child and Human Development grant (Grant ref: R01HD068437), and the National Institutes of Health (NIH, Grant ref: 5RO1AI121226-02). Genomewide genotyping data was generated by Sample Logistics and Genotyping Facilities at Wellcome Sanger Institute and LabCorp (Laboratory Corporation of America) using support from 23andMe.

**The Generation R Study**

In the Generation R Study, children were genotyped with the Illumina HumanHap 610 or 660 quad chips. A full description has been published previously (3). Data were imputed to the 1000 genomes reference panel (Phase 1 version 3). Phasing was done using MACH software, and imputation using Minimac software. DNAm was measured using the Illumina 450k array (4).

The Generation R Study is conducted by Erasmus MC, University Medical Center Rotterdam in close collaboration with the School of Law and Faculty of Social Sciences of the Erasmus University Rotterdam, the Municipal Health Service Rotterdam area, Rotterdam, the Rotterdam Homecare Foundation, Rotterdam and the Stichting Trombosedienst & Artsenlaboratorium Rijnmond (STAR-MDC), Rotterdam. We gratefully acknowledge the contribution of children and parents, general practitioners, hospitals, midwives and pharmacies in Rotterdam. The generation and management of the Illumina 450K methylation array data for the Generation R Study was executed by the Human Genotyping Facility of the Genetic Laboratory of the Department of Internal Medicine, Erasmus MC, the Netherlands. We thank Mr. Michael Verbiest, Ms. Mila Jhamai, Ms. Sarah Higgins, Mr. Marijn Verkerk and Dr. Lisette Stolk for their help in creating the EWAS database. We thank Dr. A.Teumer for his work on the quality control and normalization scripts.

The general design of the Generation R Study is made possible by financial support from the Erasmus MC, Erasmus University Rotterdam, the Netherlands Organization for Health Research and Development and the Ministry of Health, Welfare and Sport. The EWAS data were funded by a grant from the Netherlands Genomics Initiative (NGI)/Netherlands Organisation for Scientific Research (NWO) Netherlands Consortium for Healthy Aging (NCHA; project nr. 050-060-810), by funds from the Genetic Laboratory of the Department of Internal Medicine, Erasmus MC, and by a grant from the National Institute of Child and Human Development (R01HD068437). This project has received funding from the European Union’s Horizon Europe Research and Innovation Programme under grant agreement nº 101137146 (STAGE). Views and opinions expressed are however those of the author(s) only and do not necessarily reflect those of the European Union. Neither the European Union nor the granting authority can be held responsible for them.

**Drakenstein Child Health Study (DCHS)**

Genotyping was performed with the Global Screening Array. DNAm was measured using the Illumina EPIC version 2.

Consequently, a small proportion of mQTL-CpG pairs could not be replicated due to either unavailability of data or violation of the positivity assumption in certain genotype groups.

### Note S2. Definition of four genetic effect trajectory types

Time‐specific genetic effect estimates (β) for each SNP-CpG pair were calculated at ages 0, 7, 15, and 24 years. Each trajectory is characterized by a line through the four time-specific β coefficients. All β values for a pair were multiplied by −1 (flipped) if: (i) the birth effect was significantly negative (p < 0.05), or (ii) the most negative effect was significant (p < 0.05) and the birth effect was not significantly positive. Then, differences between adjacent time-specific effects were assessed with two-sided Wald tests, and each time segment was labelled as increasing, decreasing, or constant (if p ≥ 0.05). Pairs were assigned to one mutually exclusive class using the rules below:

| Category | Pattern | Meaning |
| --- | --- | --- |
| Sign-reversed | At least one significantly positive β and one significantly negative β | Direction of the genetic effect switched across development. |
| Increasing | ≥1 significant increase; no significant decrease segments | Genetic influence strengthened with age. |
| Decreasing | ≥1 significant decrease; no significant increase segments | Genetic influence attenuated with age. |
| Fluctuating | At least one significant increase and one significant decrease; not sign-reversed | Genetic effect size rose and fell over time. |

### Note S3. Enrichment analysis for longitudinal mQTLs

Before performing analysis, effect and non-effect alleles were aligned between our study and GWAS summary statistics for the traits of interest. When SNPs were associated with multiple CpGs in our dataset, only the most significant association was retained to avoid redundancy.

For the 194,664 SNPs analyzed in this study, we first computed the Pearson’s correlation ($r$) between our estimated genotype-by-age interaction effects (β) and SNP-trait association estimates. We then applied the following binomial regression model under two complementary settings: (i) testing whether the magnitude of SNP–trait associations predicts the likelihood of a SNP being a longitudinal mQTL, and (ii) testing whether stronger genotype-by-age interaction effects predict trait associations. In the first setting, $y_{i}$ is a binary indicator denoting whether SNP $i$ exhibited a significant dynamic influence on DNAm, and $\left| z_{i} \right|$ is the absolute z-score of the SNP-trait association. In the second setting, $y_{i}$ indicates whether SNP $i$ was associated with the trait (p < 1× 10^−5^), and $\left| z_{i} \right|$ reflects the interaction effect size. The model was adjusted for linkage disequilibrium (LD) score and minor allele frequency (MAF).

$${logit(Pr(y}_{i}=1))= \beta_{0}+\beta_{1}\left| z_{i} \right|+\beta_{2}{LD}_{i}+\beta_{3}{MAF}_{i}+\epsilon_{i}$$

across $i$ where $\epsilon_{i} \sim N\left( 0,\sigma_{\epsilon}^{2} \right)$

1.	Relton CL, Gaunt T, McArdle W, Ho K, Duggirala A, Shihab H, et al. Data Resource Profile: Accessible Resource for Integrated Epigenomic Studies (ARIES). Int J Epidemiol. 2015 Aug;44(4):1181–90.

2.	Bibikova M, Barnes B, Tsan C, Ho V, Klotzle B, Le JM, et al. High density DNA methylation array with single CpG site resolution. Genomics. 2011 Oct;98(4):288–95.

3.	Medina-Gomez C, Felix JF, Estrada K, Peters MJ, Herrera L, Kruithof CJ, et al. Challenges in conducting genome-wide association studies in highly admixed multi-ethnic populations: the Generation R Study. Eur J Epidemiol. 2015 Apr;30(4):317–30.

4.	Lehne B, Drong AW, Loh M, Zhang W, Scott WR, Tan ST, et al. A coherent approach for analysis of the Illumina HumanMethylation450 BeadChip improves data quality and performance in epigenome-wide association studies. Genome Biol. 2015 Feb 15;16(1):37.

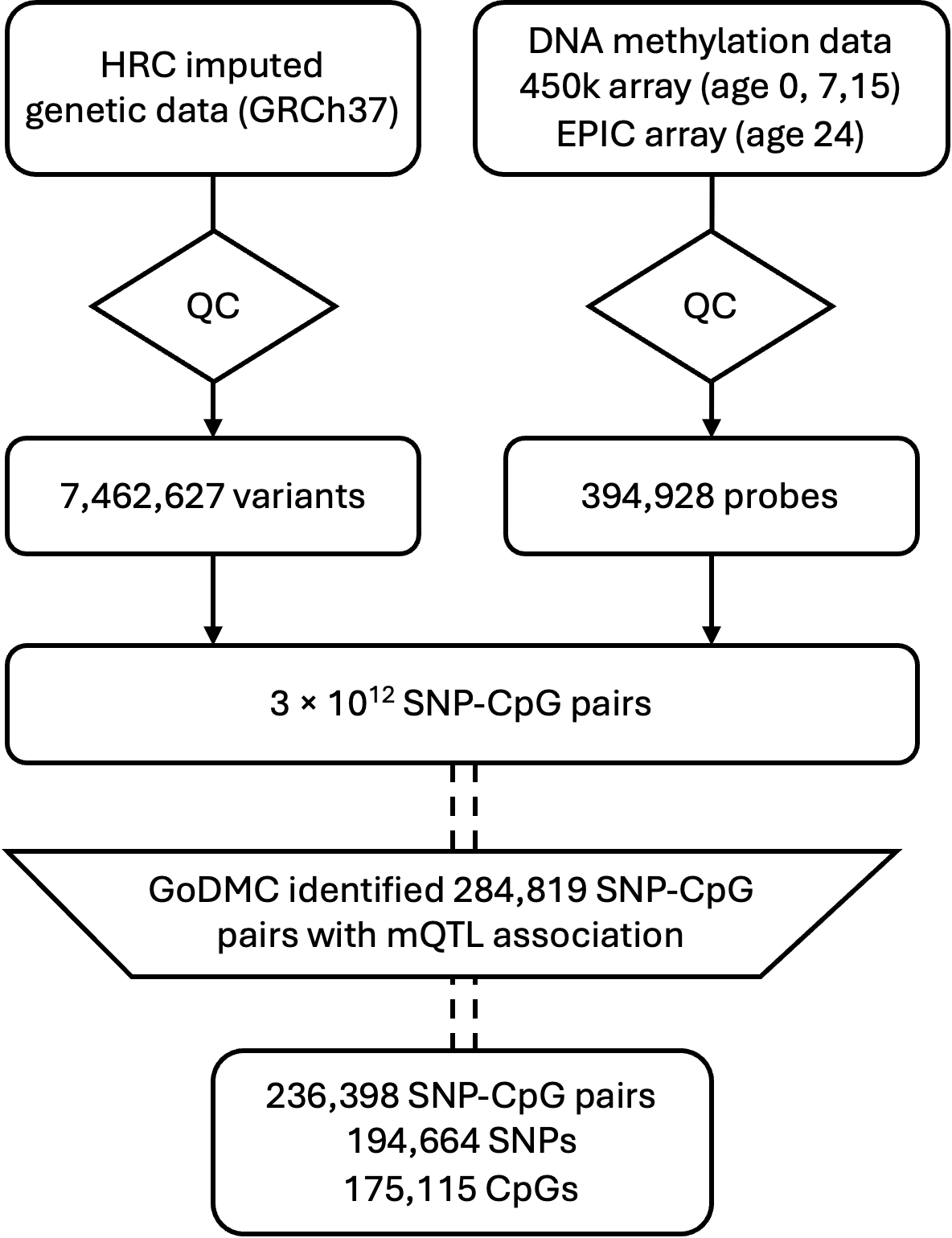

### Figure S1. Workflow for selecting the analysis set of SNP-CpG pairs

HRC = Haplotype Reference Consortium, QC = quality control, GoDMC = Genetics of DNA Methylation Consortium, mQTL = methylation quantitative trait locus.

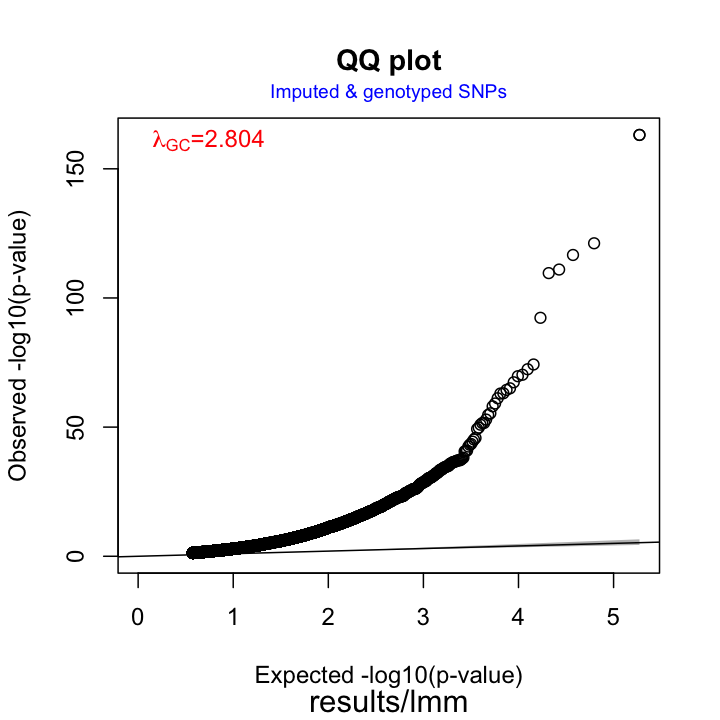

### Figure S2. Q-Q plot of the LMM assessing the genotype-age interaction effects on DNAm

Observed and expected *p*-values are on a –log_10_ scale. The black diagonal line represents the null hypothesis of no association, and deviations above the line indicate an excess of small *p*-values. LMM = linear mixed model.

a

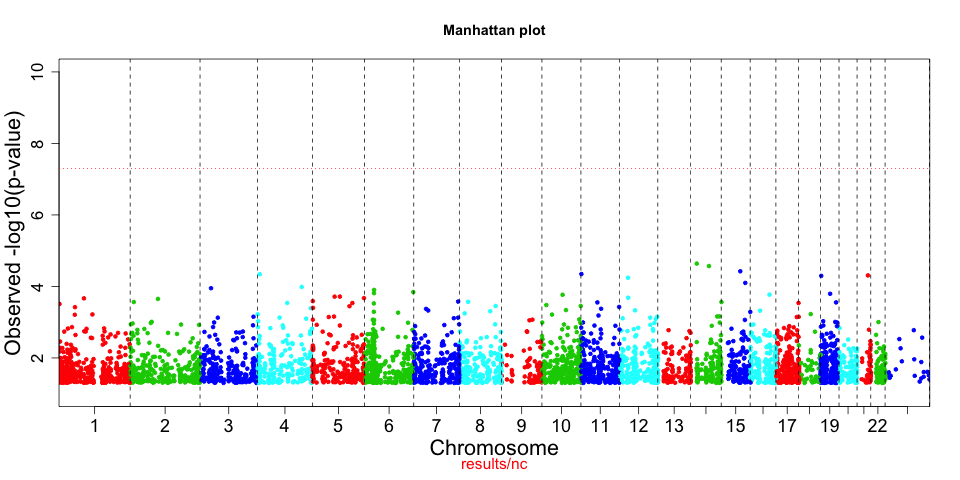

b

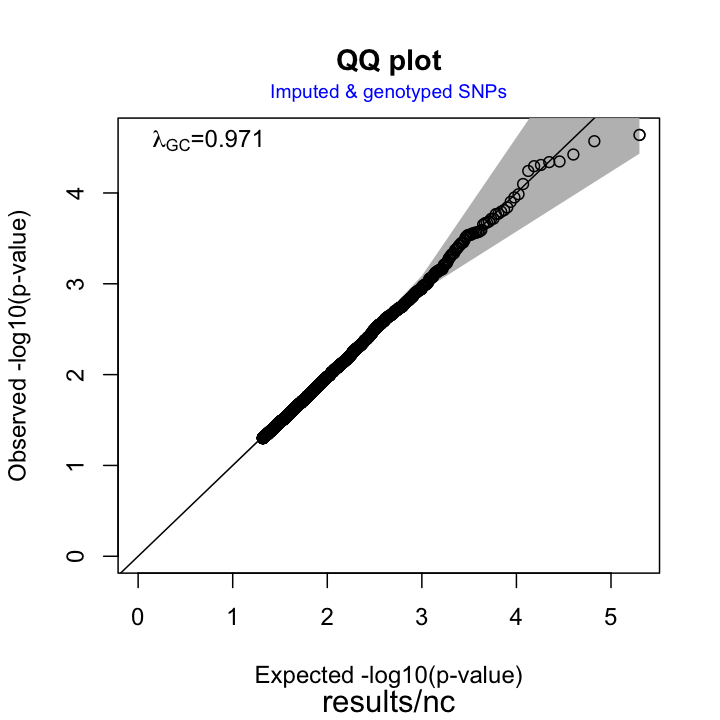

### Figure S3. Manhattan plot (a) and Q-Q plot (b) for the negative control analysis

Plots are based on the genotype-age interaction term from the LMM, applied to 100,000 SNP-CpG pairs expected to show no association. The model settings were identical to those used in the main analysis. The red dashed line corresponds to the genome-wide significance threshold of p = 5 × 10^−8^. *P*-values are on a –log_10_ scale. LMM = linear mixed model.

a	b

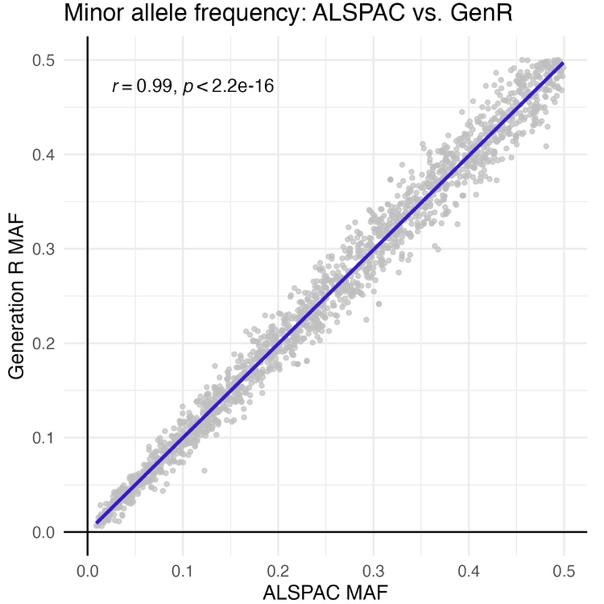

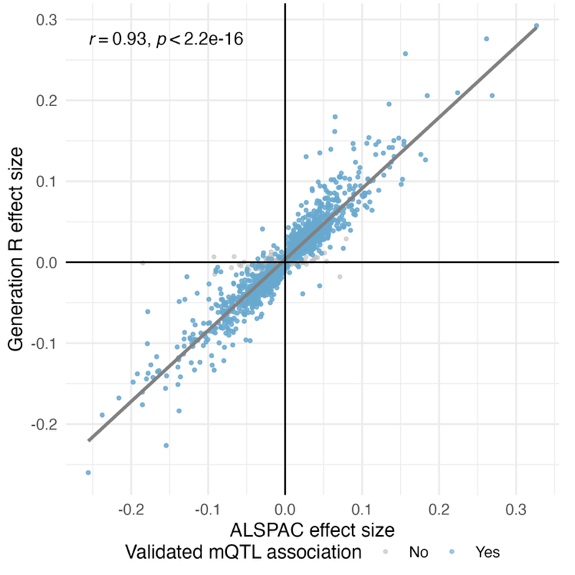

c	d
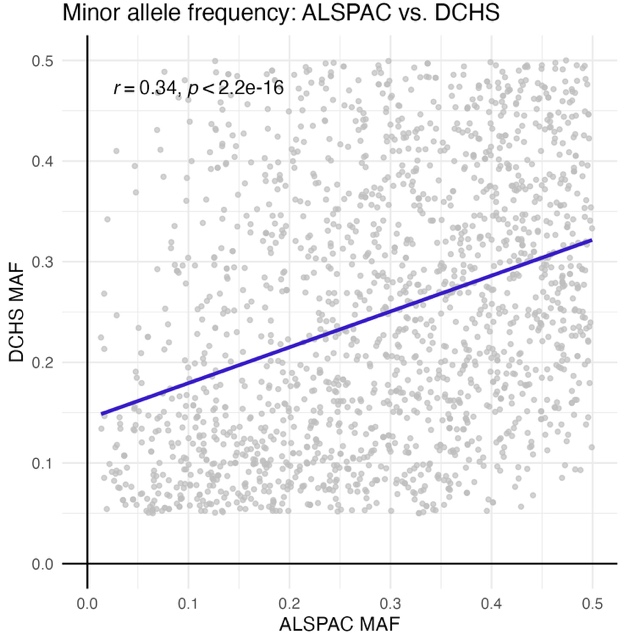

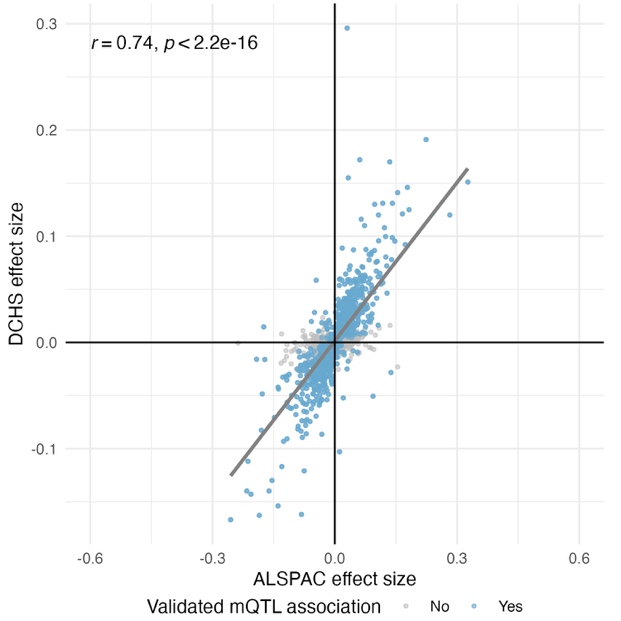

### Figure S4. Replication of genetic main effect for identified longitudinal mQTLs.

The Pearson’s correlation coefficient (r) quantifies concordance across studies. **a** Comparison of SNP MAFs between the discovery cohort (ALSPAC) and the Generation R Study. **b** Comparison of SNP main effects between ALSPAC and the Generation R Study. Points are color-coded according to whether SNP-CpG pairs showed significant mQTL associations in the replication cohorts based on linear models. Pairs denoted by blue dots passed QC and were further tested for genotype-by-age interaction using LMMs. **c** Comparison of SNP MAFs between the discovery cohort (ALSPAC) and the DCHS cohort. **d** Comparison of SNP main effects between ALSPAC and the DCHS cohort. MAF = minor allele frequency, QC = quality control, LMM = linear mixed model, ALSPAC = Avon Longitudinal Study of Parents and Children, DCHS = Drakenstein Child Health Study.

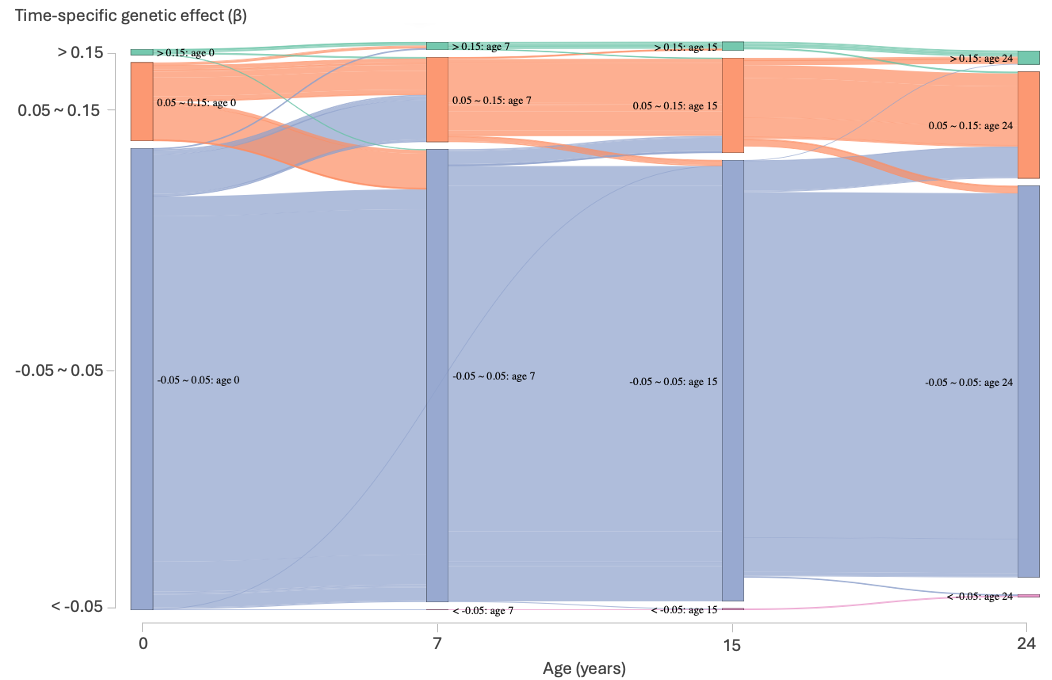

### Figure S5. Sankey plot illustrating the magnitude of longitudinal mQTL–CpG associations

Each stream in the Sankey plot represents a group of identified SNP-CpG pairs categorized by the magnitude of their associations at each time point. Colours correspond to effect size categories. Widths of the flows indicate the number of pairs falling within each effect size category, allowing visualization of how the strength of genetic effect changes over time.

a

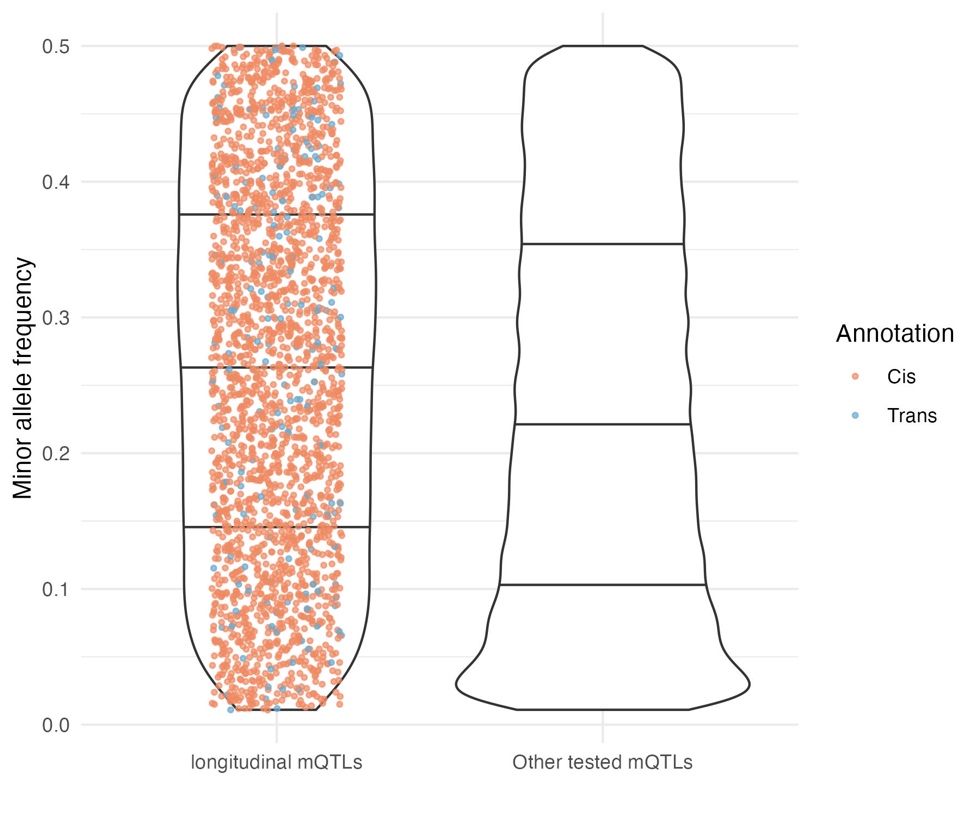

b

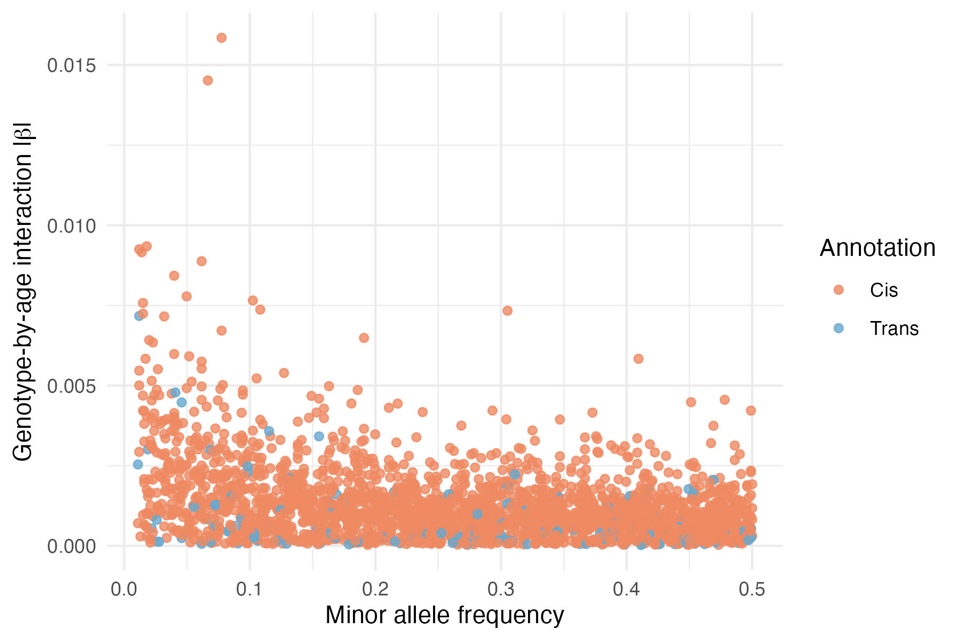

### Figure S6. Minor allele frequency distribution of longitudinal mQTLs

Points are color-coded according to the cis/trans positional classification of longitudinal mQTLs. **a** Minor allele frequency distribution (y-axis) of identified longitudinal mQTLs versus other tested mQTLs. The width of each violin indicates the local density of SNPs across the allele frequency spectrum. Black solid lines represent the 25th, 50th, and 75th percentiles. **b** Minor allele frequency (x-axis) versus absolute effect size (y-axis) for identified longitudinal mQTLs.

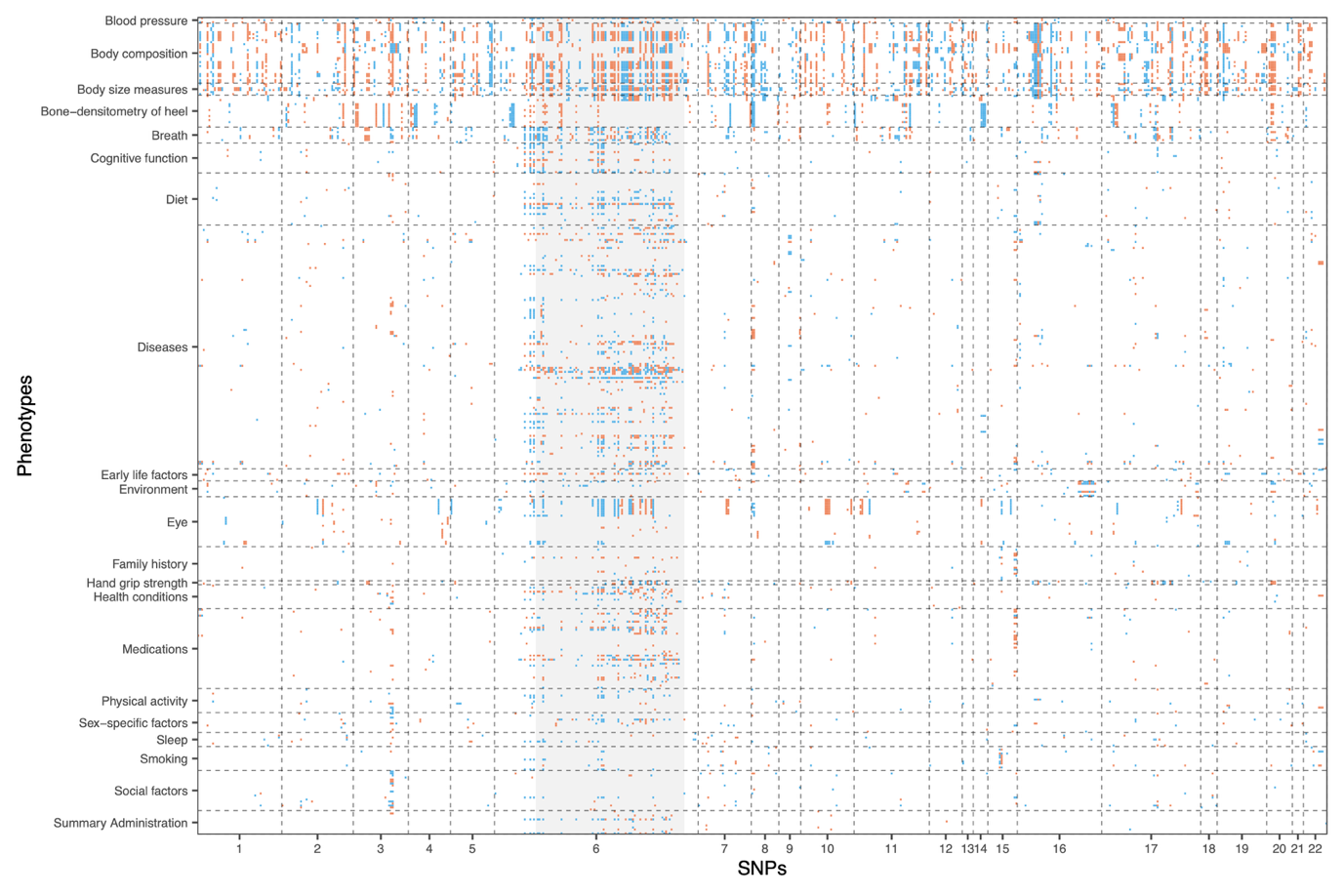

### Figure S7. Overlap of associations between longitudinal mQTLs and UK Biobank phenotypes.

PheWAS scan was conducted using summary statistics from previously published GWAS available in the OpenGWAS database. A *p*-value threshold of 1× 10^−5^ was used to define significance. To avoid overcrowding the heatmap, we only displayed results involving the 616 SNPs associated with more than one phenotype and the 409 phenotypes associated with more than one SNP here. Each cell represents a SNP-phenotype association, with red and blue indicating positive and negative effect directions, respectively. Phenotypes on the y-axis are grouped to higher-level categories to facilitate interpretation, while longitudinal mQTLs on the x-axis are ordered by chromosomal position. The grey-shaded region denotes the HLA region (chr 6: 28477797-33448354). PheWAS = phenome-wide association study, GWAS = genome-wide association study, SNP = single nucleotide polymorphism, HLA = Human Leukocyte Antigen.

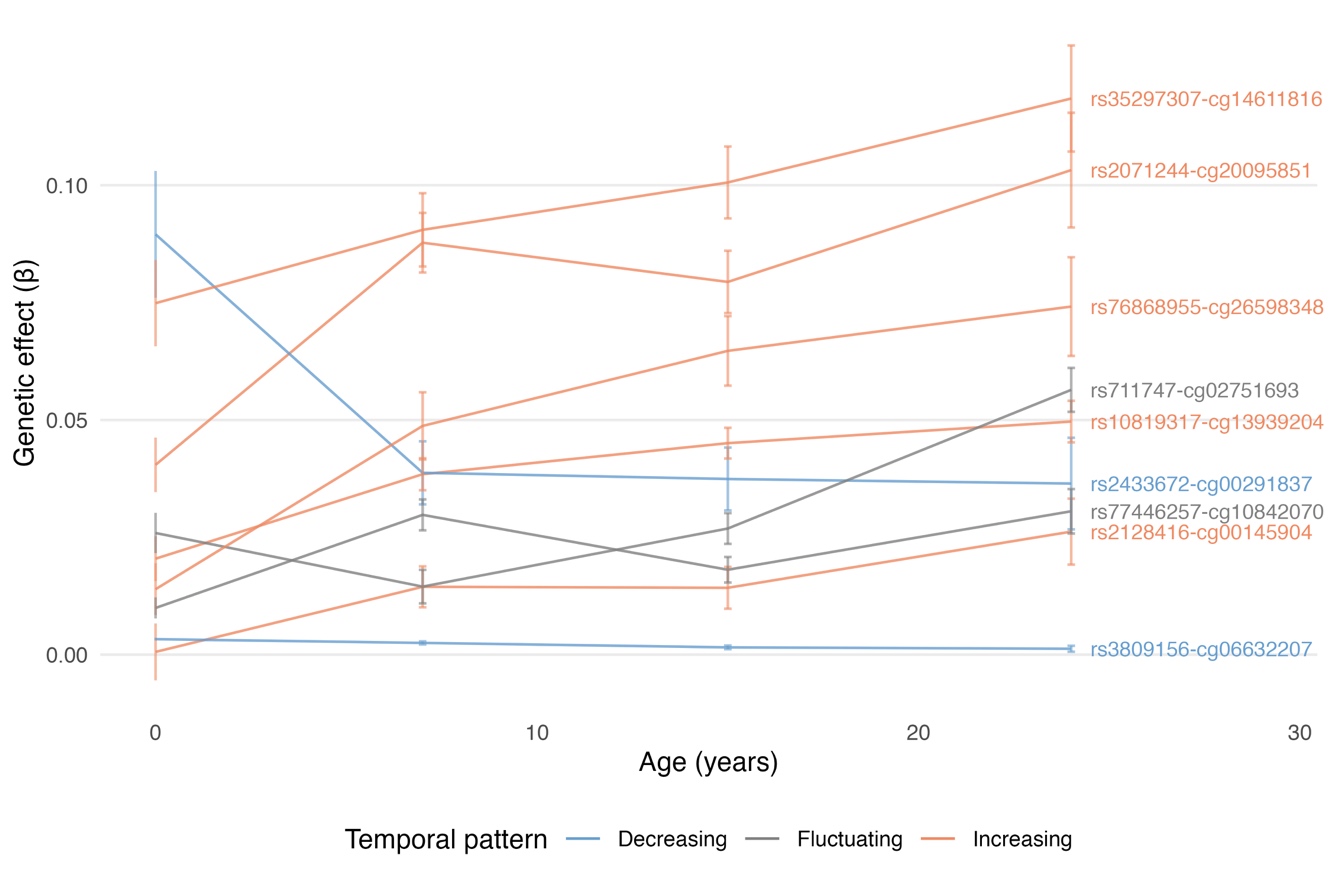

### Figure S8. Association trajectories of 9 longitudinal mQTL-CpG pairs overlapping with previously reported cell type imeQTL-CpG pairs.

Each trajectory represents β coefficients (time-specific genetic effects on DNAm) with corresponding error bars at 4 time points. Curves are color-coded according to the temporal patterns of genetic influence. imeQTL= interaction methylation QTL.

a

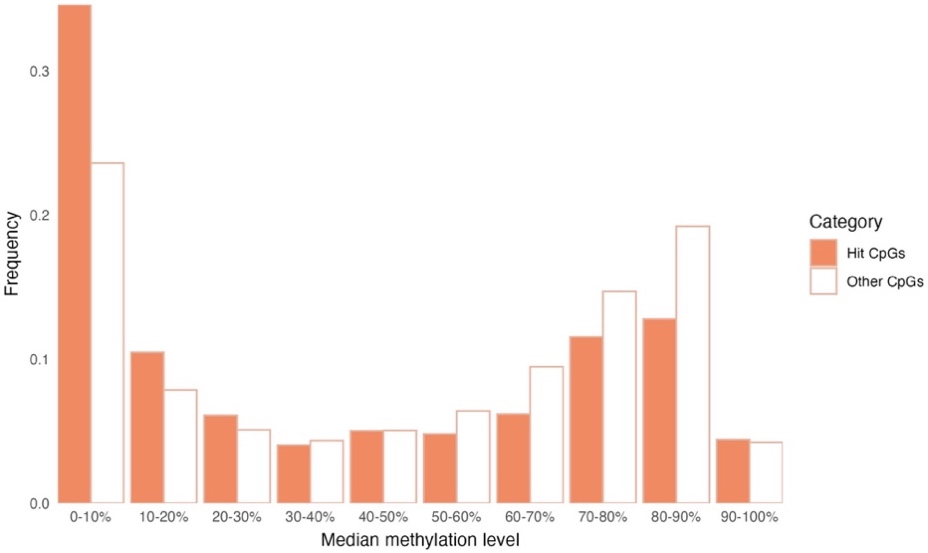

b

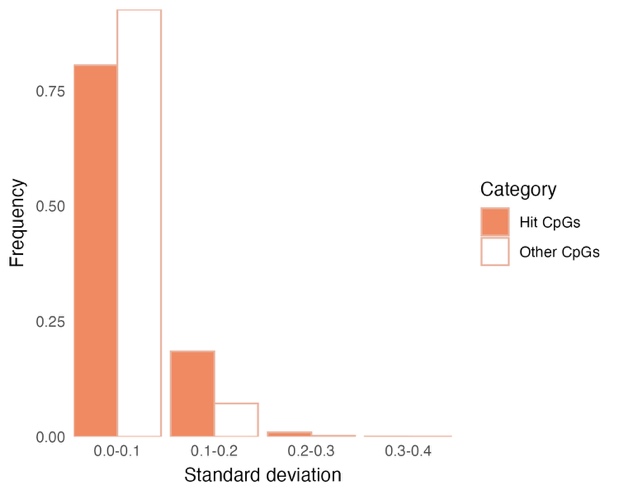

c

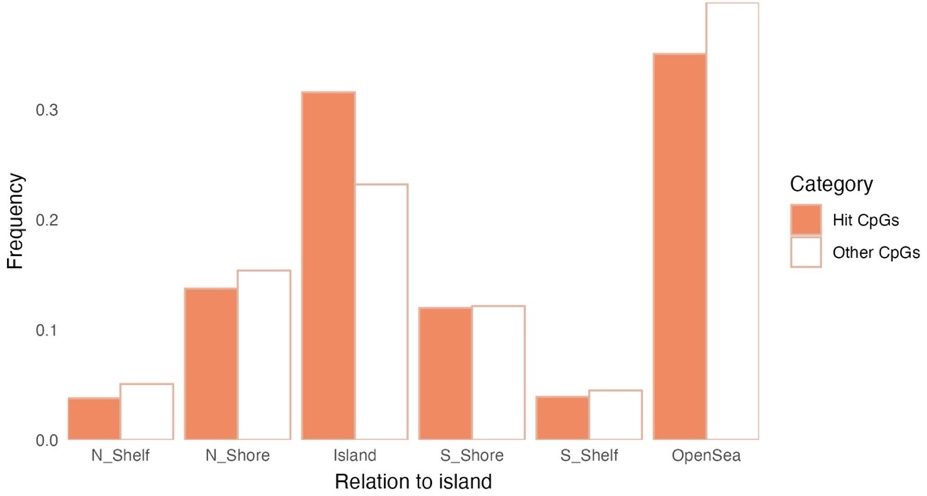

### Figure S9. Characteristics of CpG sites involved in longitudinal mQTL associations

Comparisons were made between CpG sites associated with longitudinal mQTLs and all other tested CpG sites. To calculate the statistics, DNAm values were pooled across all four time points. **a** Distribution of median DNAm levels. **b** Distribution of the standard deviation of DNAm levels. **c** Genomic distribution of CpG sites relative to CpG islands.

**a**

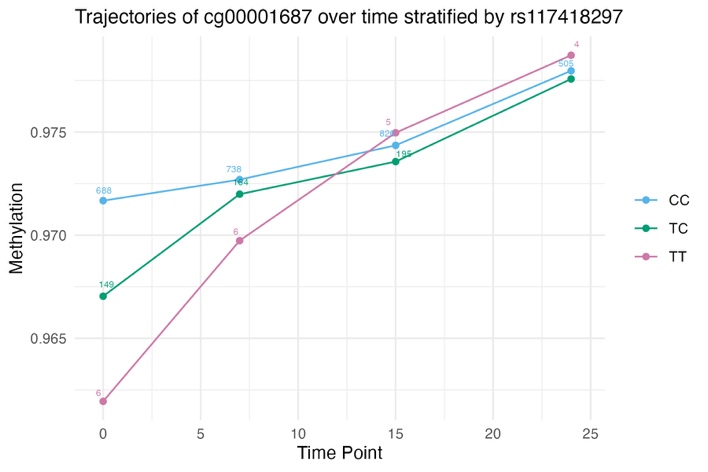

**b**

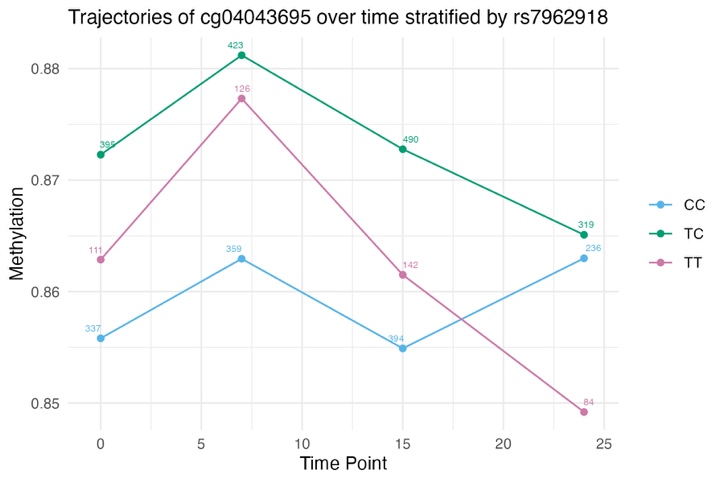

**c**

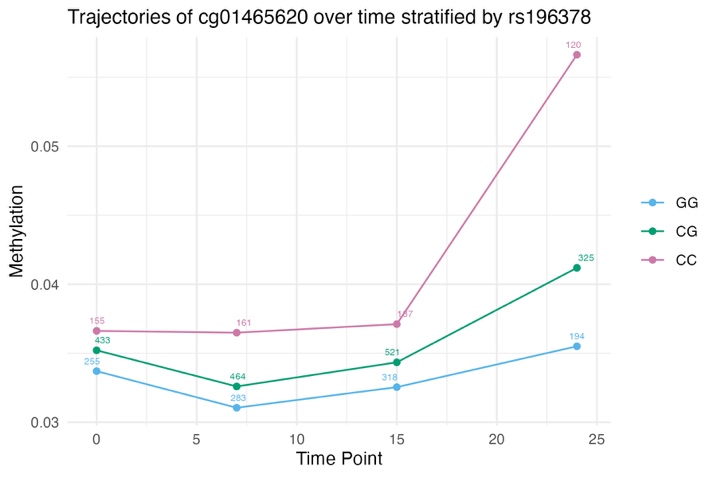

### Figure S10. DNAm trajectories for 3 longitudinal mQTL-CpG pairs

Mean DNAm levels are plotted across four time points, stratified by genotype, illustrating genotype-dependent changes in CpG methylation over time. **a** Trajectories for the rs117418297-cg00001687 pair. **b** Trajectories for the rs7962918-cg04043695 pair. **c** Trajectories for the rs196378-cg01465620 pair.

a

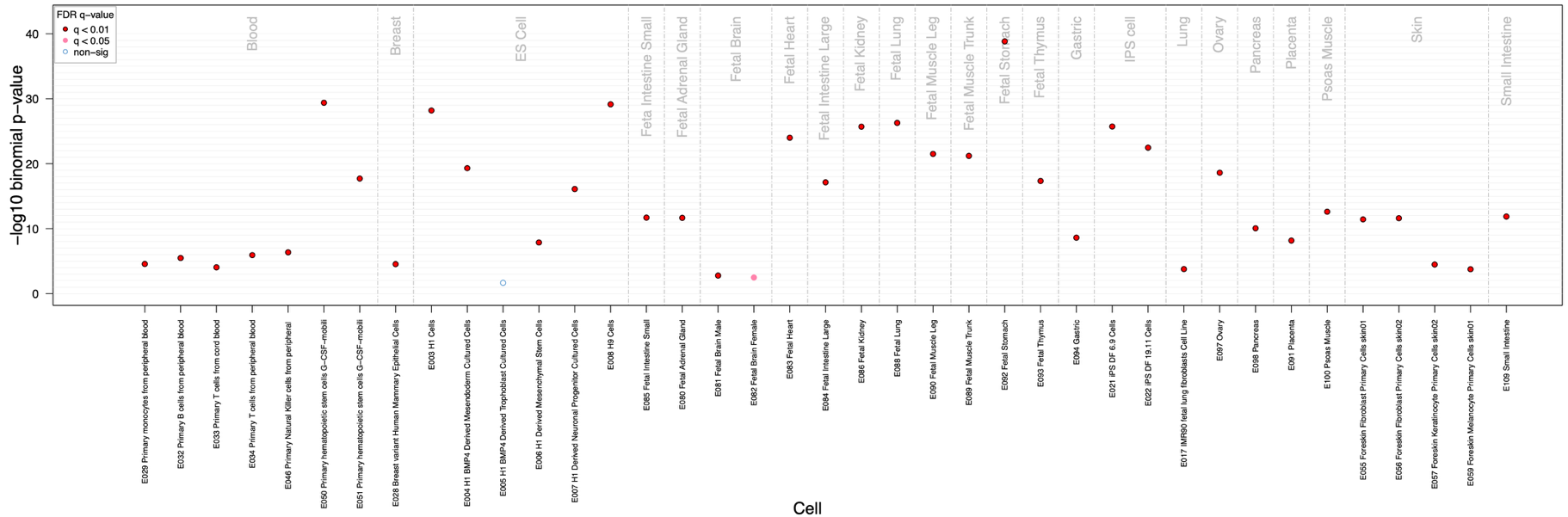

b

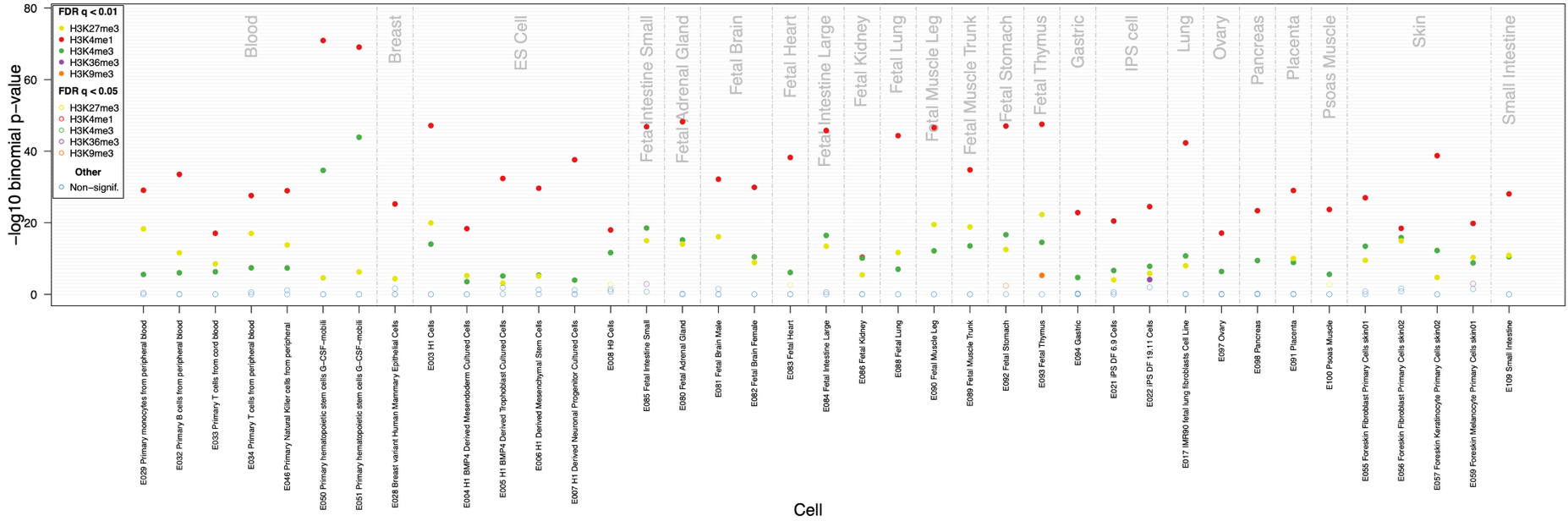

c

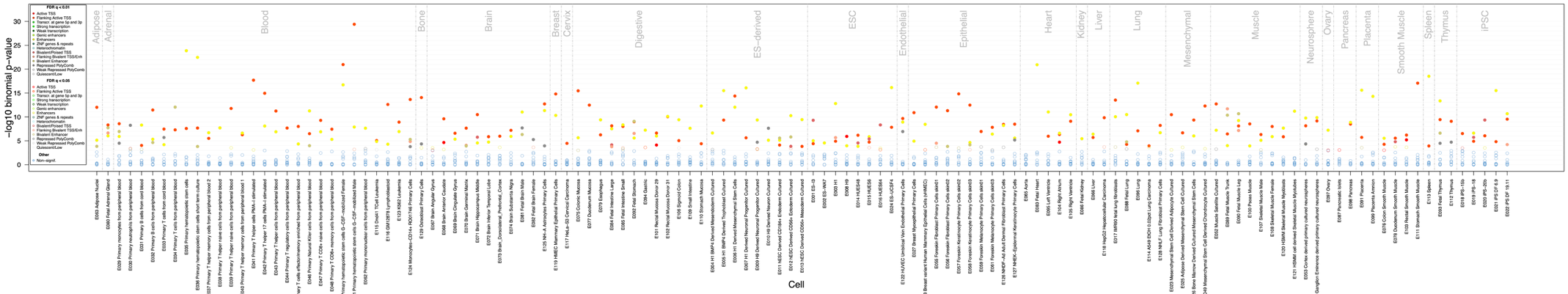

### Figure S11. Enrichment of 2329 identified CpGs in epigenetic tracks

Enrichment analysis was conducted using eFORGE v2.0 with 1,000 background repetitions and default parameters. All epigenetic annotations were obtained from the Roadmap Epigenomics Consortium. The x-axis denotes different cell types, and the y-axis represents the –log₁₀ transformed p-values, corrected using the Benjamini–Yekutieli approach (a false discovery rate approach accounting for the non-independence between epigenomic datasets). **a** Enrichment in DNase I hypersensitive sites based on the 2015 Roadmap DNase-seq hotspot dataset. DNase I hypersensitive sites indicate open chromatin regions permissive to gene expression. **b** Enrichment in 5 histone modification marks. H3K4me1, H3K4me3, and H3K27me3 mark enhancer regions, promoter regions, and reversible gene repression states involved in developmental gene silencing, respectively. **c** Enrichment in 15 chromatin states.

a

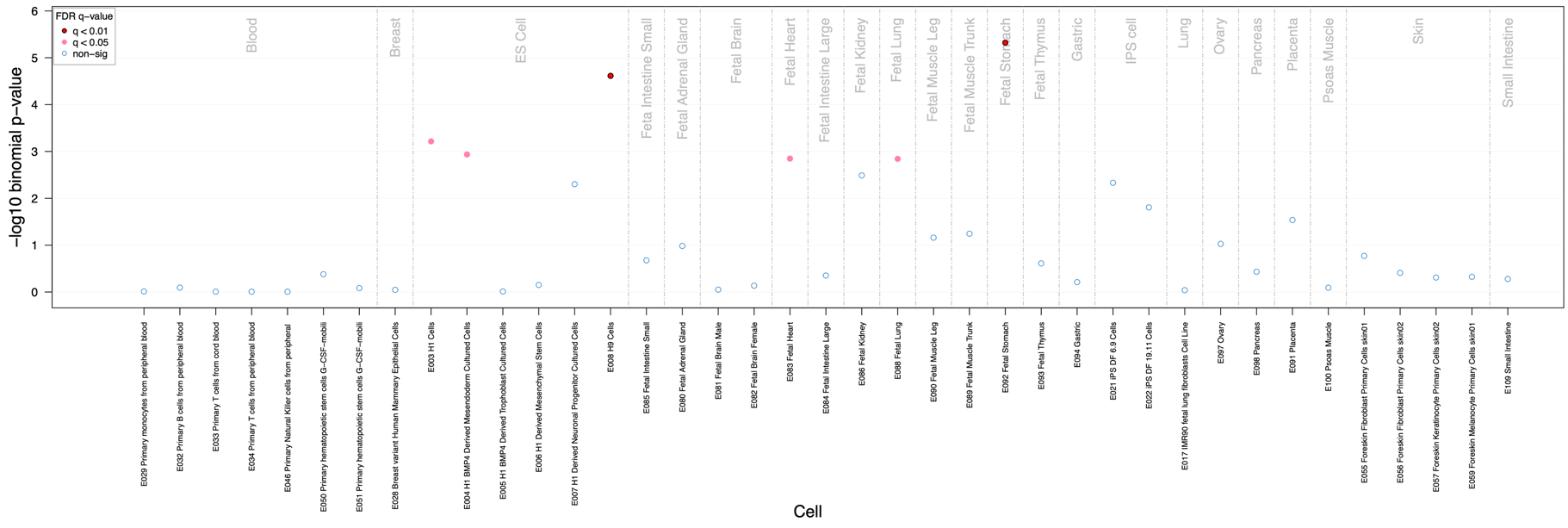

b

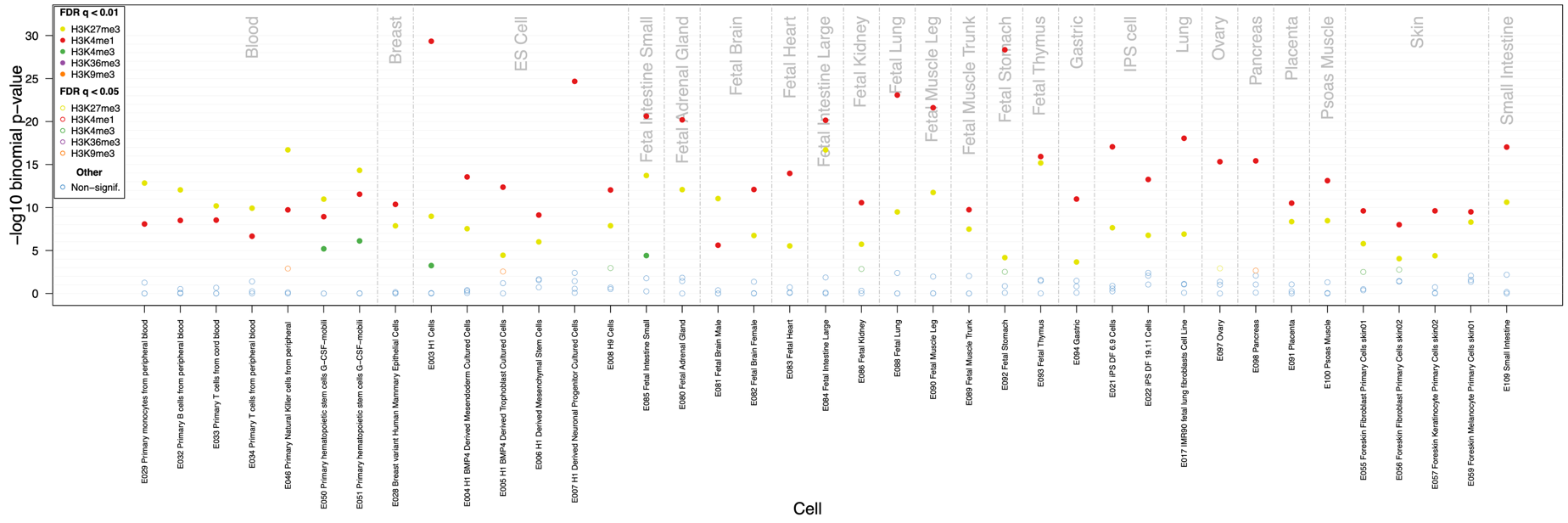

c

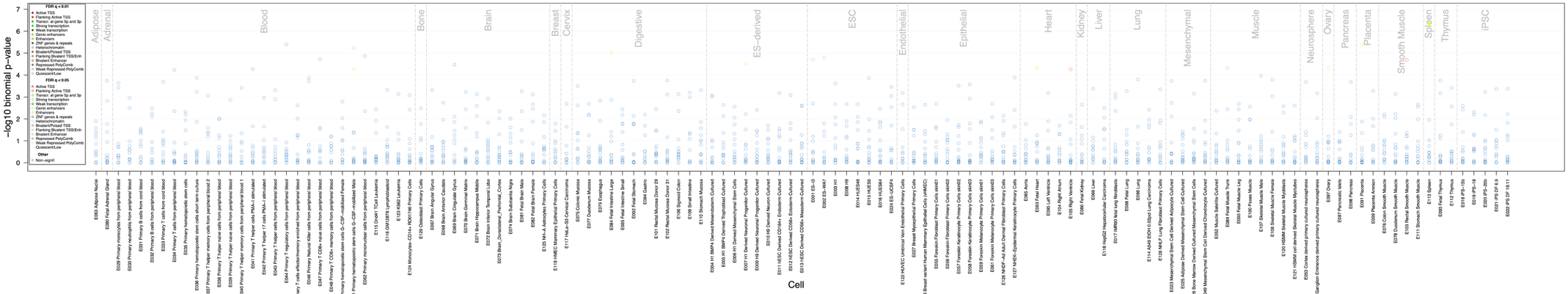

### Figure S12. Enrichment of 2329 genotype-associated CpGs in epigenetic tracks

This enrichment analysis was performed on a randomly selected set of CpGs drawn from significant signals reported by the GoDMC study, serving as a reference for comparison. The analysis settings were identical to those used for the enrichment of longitudinal mQTL associated CpGs. The x-axis denotes different cell types, and the y-axis represents the –log₁₀ transformed p-values, corrected using the Benjamini–Yekutieli approach. **a** Enrichment in DNase I hypersensitive sites based on the 2015 Roadmap DNase-seq hotspot dataset. **b** Enrichment in 5 histone modification marks. **c** Enrichment in 15 chromatin states. GoDMC = Genetics of DNA Methylation Consortium.

a

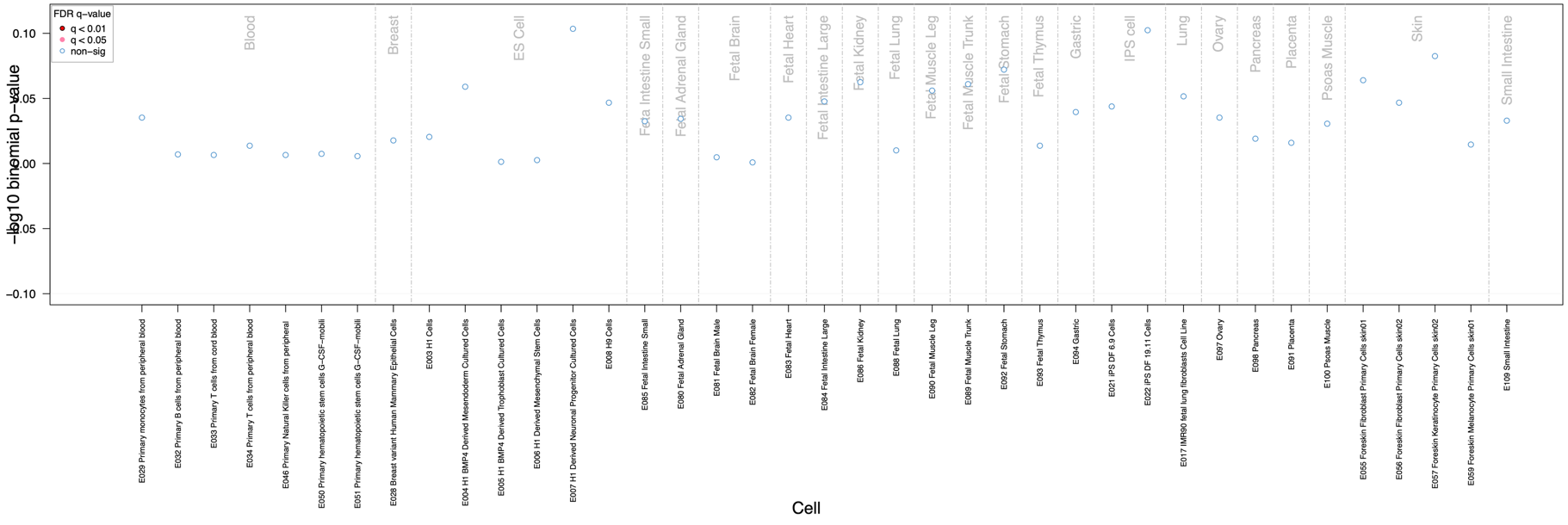

b

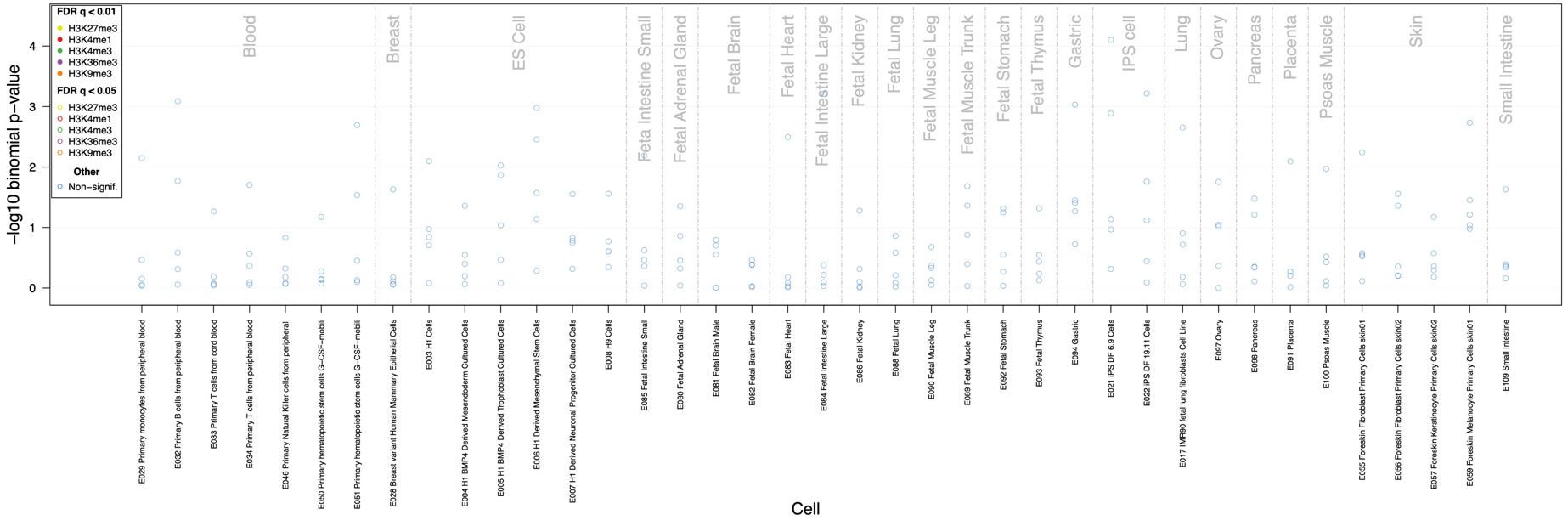

c

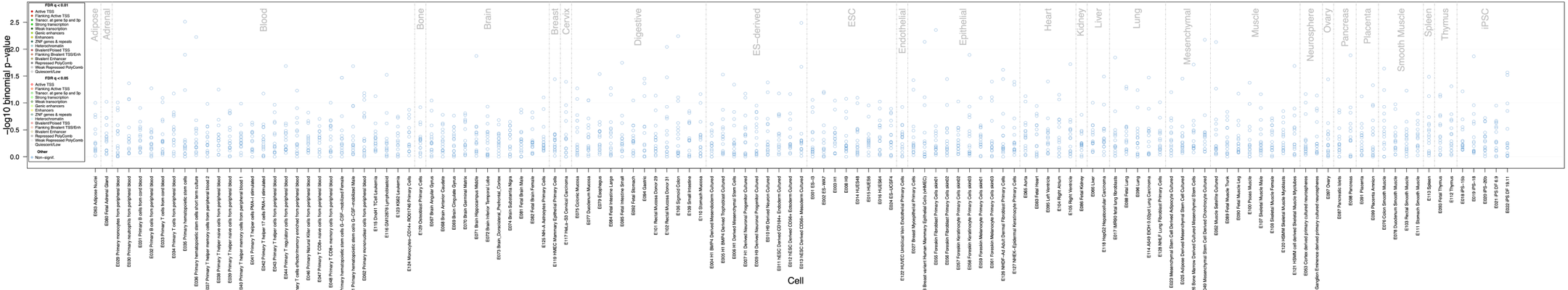

### Figure S13. Enrichment of 2329 time-varying CpGs in epigenetic tracks

This enrichment analysis was performed on a randomly selected set of CpGs drawn from significant signals reported by a longitudinal EWAS (Mulder et al., 2021), serving as a reference for comparison. The analysis settings were identical to those used for the enrichment of longitudinal mQTL associated CpGs. The x-axis denotes different cell types, and the y-axis represents the –log₁₀ transformed p-values, corrected using the Benjamini–Yekutieli approach. **a** Enrichment in DNase I hypersensitive sites based on the 2015 Roadmap DNase-seq hotspot dataset. **b** Enrichment in 5 histone modification marks. **c** Enrichment in 15 chromatin states. EWAS = epigenome-wide association study.
